# Structural visualisation of the dynamics of DNA unwinding by a replicative helicase

**DOI:** 10.1101/2024.04.19.590217

**Authors:** Taha Shahid, Muhammad Tehseen, Lubna Alhudhali, Alice Clark, Christos G. Savva, Samir M. Hamdan, Alfredo De Biasio

## Abstract

Ring-shaped hexameric helicases are nucleotide hydrolases that unwind double-stranded DNA into single strands, a necessary step in DNA replication. Two main questions are critical to understanding their function: the location of DNA strand separation within the helicase, and the precise dynamic linkage between nucleotide hydrolysis and DNA translocation. We explore these questions by employing cryo-EM to visualize the translocation mechanics of a AAA+ helicase, the SV40 Large Tumor Antigen (LTag), on forked DNA. We find the DNA fork nexus is positioned deep within the core helicase domain, with each DNA-binding loop from the six subunits securing the tracking strand by forming a paired staircase spiral that substitutes the passive strand. This structure forges an internal separation wedge, channeling the passive strand through a gap in the subunit C-tiers at the back of the helicase. Via cryo-EM continuous heterogeneity analysis we capture, and exhaustively model, a seamless spectrum of conformations of translocating LTag at high resolution. ATP hydrolysis at the tightest inter-subunit interface operates like an ‘entropy switch’, triggering coordinated rigid-body rotations of the subunit C-tiers and DNA-binding loops, resulting in directional escorting of the tracking strand through the central channel, concomitantly setting preparatory steps for cycle restart. Overall, we demonstrate a dynamic model for hydrolysis-coupled translocation and DNA unwinding by a model helicase active in replication, with implications for origin unwinding. High structural conservation of core helicase regions suggests this mechanism is applicable to hexameric helicases across domains.

## Main

### Genome unwinding is an essential process facilitated by ring-shaped hexameric helicases, highly conserved enzymes that play critical roles in DNA replication and also contribute to DNA recombination and repair ^1,2^

The Simian Virus 40 (SV40) Large Tumour Antigen (LTag) is a AAA+ helicase that facilitates both initiation and progression of viral replication^3–5^, paralleling the roles of eukaryotic CDC45-MCM-GINS (CMG) ^6–8^. LTag’s bipartite architecture comprises an N-terminal origin binding domain (OBD), and a C-terminal helicase domain with an ATPase motor for translocation ^9–12^. To initiate replication, helicase subunits assemble as head-to-head hexamers at the origin, causing local duplex melting ^13–17^. The two hexamers then commence hydrolysis-driven DNA unwinding, whereby each helicase pulls its respective tracking strand in opposite directions, inducing DNA shearing ^18^. This head-to-head topology necessitates that each hexamer expels one strand from its interior, enabling the helicases to move past one another and form separate replication forks ^19,20^.

Despite decades of study on LTag and other helicases, molecular details of hydrolysis-coupled DNA translocation and unwinding remain largely elusive ^1^. Current mechanistic models derive from crystal and cryo-EM structures reconstituted in inactive conditions ^10,11,21–25^; or fragmented enzymatic snapshots obtained under hydrolysing conditions ^26,27^, often at limited resolutions due to the inherent caveats of discretely classifying this fundamentally continuous process. Specifically, the site and mechanism of strand separation, the basis for consistent discrimination between tracking and paired strands and, critically, the allosteric couplings that relay nucleotide status between active sites to coordinate nucleotide hydrolysis and DNA translocation, remain poorly understood.

To answer these questions, we employ a combination of cutting edge cryo-EM methodologies, including continuous heterogeneity analysis ^28^, deep-learning based map postprocessing ^29^, and molecular dynamics flexible fitting ^30^ to resolve a seamless spectrum of translocation states of LTag, linking nucleotide processing with DNA progression through the helicase. We rigorously model a total of 15 reaction intermediates representing the concurrent processes of ATP hydrolysis, DNA translocation, ADP/ATP exchange, and interface reset in order to restart the cycle.

Together with a further four high-resolution structures of LTag in non-hydrolytic conditions with and without DNA, we present a holistic and empirically-derived model of nucleotide-dependent substrate translocation and unwinding by a replicative helicase.

We reveal that the DNA-binding loops mimic and eventually displace the paired strand from inside the helicase, providing new insights into the largely unexplained mechanism of DNA melting at the replication origin. Pivotally, we show that as opposed to directly powering translocation, ATP serves as a block to translocation by maintaining the bottom (DNA-binding/ATPase) tier in a tensioned state relative to the top (collar) tier, which is released upon hydrolysis – ratcheting the DNA downward. Thereby, we demonstrate that unlike previously thought, coupling of nucleotide state with DNA progression is achieved through a trans-allosteric effect, brought about by correlated rigid-body motions of C-relative to N-tiers as a result of hydrolytic reaction at the single hydrolysis-competent subunit interface in each cycle.

High structural conservation of core helicase regions suggests our conclusions extend to hexameric helicases as a whole, including CMG. This enables us to propose a cross-species, unified mechanism for helicase function.

### Helicase primed on fork DNA in a pre-hydrolytic state

To begin our studies of the basis of translocation and unwinding by a AAA+ helicase, we reconstituted LTag with a forked DNA substrate in the presense of ATP – but without Mg^2+^, preventing nucleotide hydrolysis. Thereby, we obtained a cryo-EM reconstruction of this complex at a resolution of 3.1 Å (Figure 1a; Extended Data Figure 1; Extended Data Table 1). The helicase core is structured as a two-tiered hexameric ring. Its six oligomerization domains (N-tiers) form a collar, with the bulkier ATPase motor domains (C-tiers) lying beneath. The OBD and J-domains exhibit significant flexibility, as indicated by fuzzy density in 2D class averages (Extended Data Figure 1). The DNA fork-junction is nestled deep inside the central channel within the collar domain, with the tracking strand held firmly by the six DNA binding loops in the C-tier chamber, while the unpaired passive strand is flexible and disordered, and hence invisible (Figure 1b-c). The N-tiers slightly deviate from six-fold rotational symmetry. However, a significant departure from this symmetry is observed amongst C-tiers. This lies in contrast to the “apo” configuration of ATP-LTag derived from DNA-absent particles (Extended Data Figure 2; Extended Data Table 1), which mirrors the C6 symmetry previously identified in crystallographic analyses ^10,11^ (Extended Data Figure 3). Asymmetry in the DNA-bound form arises from systematic histidine-nucleobase and lysine-phosphate (as well as arginine-phosphate) engagements of the internal DNA binding domains with the tracking strand (Figure 1c). These interactions drive progressive rotations of the C-tiers, fostering the unique staircase configuration of the DBDs in alignment with the DNA backbone (Extended Data Figure 3). In both DNA-anchored and apo forms, each subunit associates with an ATP molecule. However, in the apo form the DBDs appear equiplanar and partly flexible (Extended Data Figure 3).

**Figure 1.**
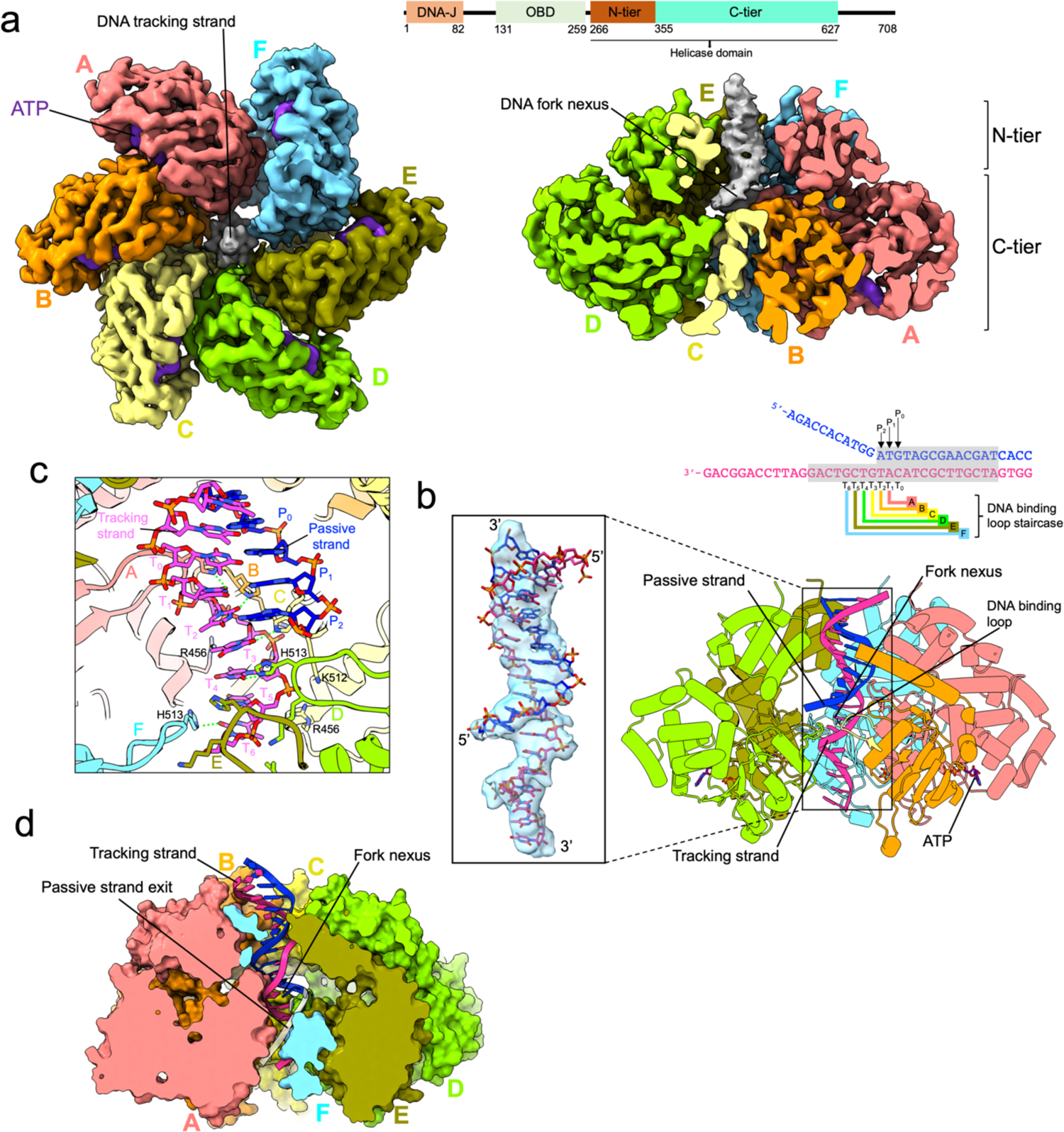
Cryo-EM structure of the LTag hexamer bound to forked DNA and ATP in a pre-hydrolysis state. **a** Bottom and side views of the cryo-EM reconstruction of the LTag hexamer bound to forked DNA and ATP. In the side view, the map has been sliced to reveal the DNA density crossing the N-tier. Above, the domain structure of the LTag monomer is illustrated. **b** Molecular model in ribbon representation. The model has been sliced to highlight the DNA binding loops staircase intruding into the DNA minor groove at the fork nexus. The inset shows the map density around the DNA model in stick representation. The DNA substrate sequence is shown, with the modelled portion shaded in gray. **c** Molecular interactions at the DNA nexus. Hydrogen bonds between H513 and nucleobases are shown as green dotted lines. Below the P2 base of the passive strand, the DNA binding loops of the D, E, and F subunits create an internal DNA separation wedge. **d** LTag model depicted with a surface representation that has been sectioned to show its internal structure. The tracking and passive strands are displayed as sticks in pink and blue, respectively. The 5’ tail of the passive strand, which is not visible in the map, has been theoretically modeled to pass through the gap between the C-tiers of the A and F subunits and is highlighted in white.

At the fork junction, the density map shows clear definition for DNA nucleotides over several base pairs. Beyond these base pairs, density is more limited to nucleobases (Figure 1b), suggesting the DNA segment passing through the N-tier collar is somewhat flexible. A conformational distinction is also observed between the DNA segments. While the duplex region conforms to B-form DNA, the bound tracking strand is compacted with a reduced helical diameter of ∼16 Å. Sidechains of the six H513 residues follow the right-handed helical pattern of the tracking strand, establishing hydrogen bonds with successive bases. The tracking strand is further anchored by salt bridges between K512 and R456 and the backbone phosphates, the latter being positioned two nucleotides further down the staircase (Figure 1c). Due to mixed composition of the DNA substrate (Figure 1b), tracking and passive strand bases were assigned as thymidines and adenines, respectively. Nonetheless, the aromatic rings of pyrimidines and purines would both present a polar atom (O or N) at equivalent locations, available to interact with H513’s sidechain protonated nitrogen at the ε (NE2) position. At the nexus, imidazole rings of H513 belonging to the top three DNA-binding loops from the staircase intrude into the DNA minor groove. At this point, nucleobases T1 and T2 from the tracking strand not only interact with H513 but also establish Watson-Crick bonds with their complementary bases P1 and P2 on the passive strand (Figure 1c). Yet, the third base (T3) from the top may not pair with its passive strand counterpart due to interference from nearby loop residues. From this position downward, the DNA binding loops essentially occupy the space where the passive stand would otherwise be located, creating an internal separation wedge. This suggests that the passive strand, though not directly observable in the map, exits at the back through the opening between the C-tiers of the top and bottom subunits (Figure 1d). As discussed later, internal strand melting and extrusion through the C-tiers is also supported by a previously determined structure of an LTag dimer bound to origin DNA ^9^.

### Nucleotide binding and intersubunit interactions

Unlike the apo form of the helicase, whereby inter-subunit interfaces appear homogenous, the DNA-bound variant in the pre-hydrolytic state presents interfaces with varying features, exhibiting unique nucleotide coordination patterns corresponding to the specific placements of the associated DNA binding loops on the staircase (Figure 2a-b). Past crystallographic studies have revealed that conformation of most LTag *cis*-residues (those from the same monomer) which interact with ATP is maintained when bound to ADP ^10^. This observation led to the identification of critical *trans*-residues (those from the subsequent monomer); i.e., K418, R498, and R540 – often dubbed “fingers”. These residues, critical for ATP coordination and hydrolysis, serve as proxies for determining interface type. As such, the two top interfaces (A/B and B/C) in our structure conform to an “ATP-like” profile: characterized by direct interactions of the K418 sidechain with ATP α, β and γ-phosphates, R540 with the ATP γ-phosphate, and R498 with *cis*-D474; with R498 possibly further interacting with the γ-phosphate via a water molecule ^10^ (Figure 2b). Meanwhile, the C/D interface closely resembles an ADP-configuration, marked by direct interaction between R498 and the nucleotide β-phosphate. The subsequent interfaces along the staircase (D/E and E/F) align with the ADP-configuration, yet the R498 side chain is positioned at a distance that prevents direct interaction with the nucleotide. Given the staircase architecture, the bottom-most subunit (F) does not establish a pronounced interface with its top counterpart (A) (Figure 2b). The extent of buried surface area (BSA) at the interfaces correlates with the nucleotide configuration (Figure 2b). Notably, ATP-like interfaces are markedly more compact than the ADP-like ones. We analyzed the nucleotide coordination in this structure and compared it to a 3.4 Å-resolution structure derived from LTag reconstituted with ADP and a forked DNA substrate with an extended tracking strand (Extended Data Figure 4; Extended Data Table 1). The general structural framework between the two is similar, including the arrangement of the DNA binding loops relative to DNA (Extended Data Figure 5a). Yet, there are noteworthy differences in intersubunit interactions (Extended Data Figure 5b).

**Figure 2.**
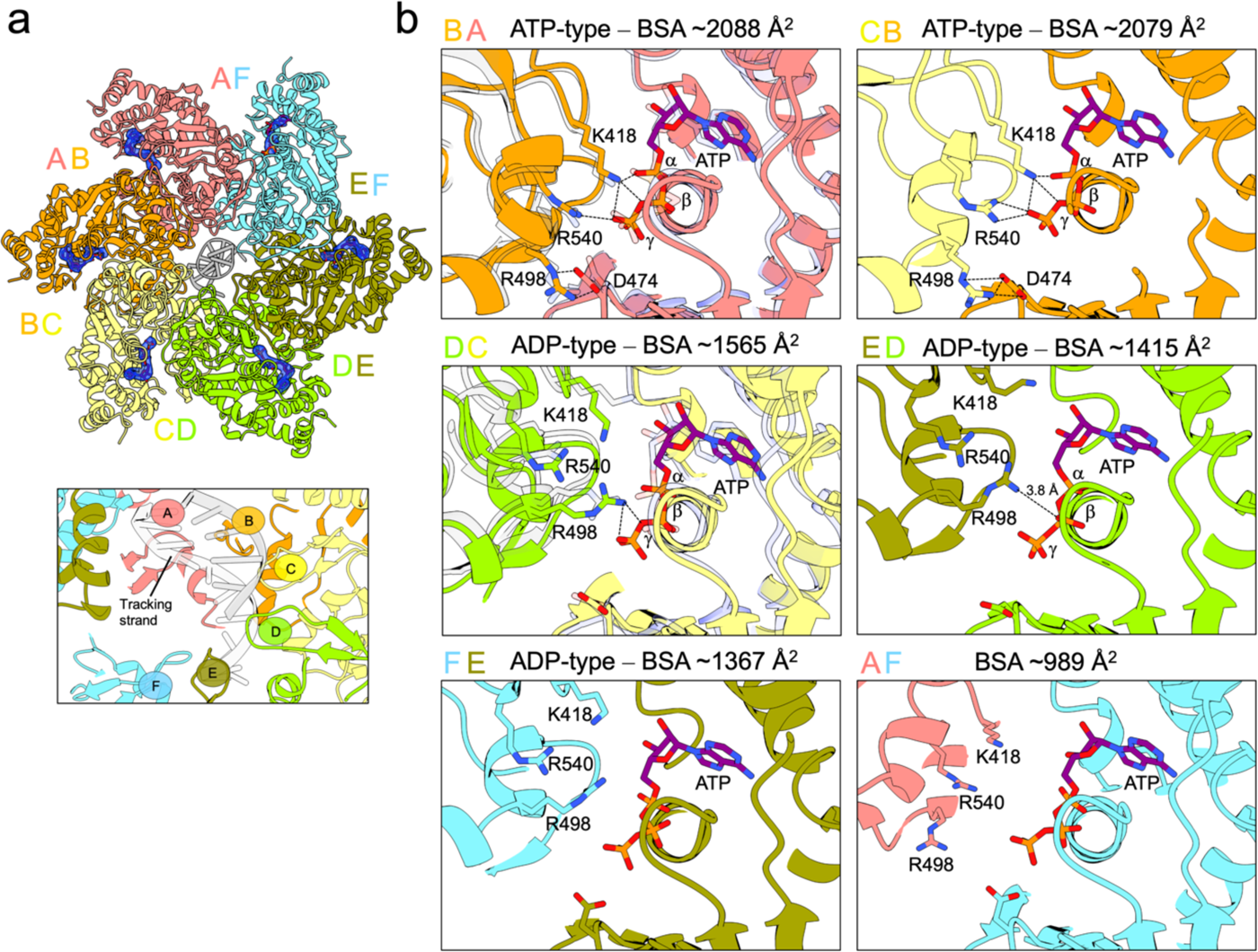
Inter-subunit interactions and ATP binding in the LTag−Forked DNA complex. **a** Bottom view of the cryo-EM model. The map density around the ATP molecules is shown in blue. The lower inset shows details of the DNA binding loop staircase and DNA. **b** Inter-subunit interactions at the nucleotide binding pocket. Calculated Buried Surface Areas (BSA) for each interface are shown, along with the assigned interface type. Hydrogen bonds are shown as black dotted lines. In the insets, chain C from the LTag-ATP crystal structure (PDB: 1SVM)^10^ is superimposed on chain A of the cryo-EM model at the AB interface, while chain B from the LTag-ADP crystal structure (PDB: 1SVL)^10^ overlays chain C at the CD interface, both shown in transparency to underscore the structural alignment.

In the ADP-bound structure, strong nucleotide coordination at the A/B and B/C interfaces is absent (Extended Data Figure 5c). As a result, the A/B and B/C interfaces are less compact than in the ATP-bound structure (Figure 2 and Extended Data Figure 5b). On the other hand, the C/D, D/E, and E/F interfaces, remain consistent between the ATP-and ADP-bound structures. This further validates the categorization of A/B and B/C interfaces as ATP-type. Interestingly, in the ADP-bound map, density at the γ-phosphate position is noticeable at the C/D, D/E and E/F interfaces (Extended Data Figure 5c). This suggests that a chloride ion may be stabilized at the γ-phosphate position by the influence of the trans-R498 finger, a feature previously documented in the E1−DNA structure bound to ADP ^25^.

We further characterized the helicase bound to forked DNA with the extended tracking strand, and a non-hydrolysable ATP analog (AMP-PNP), in the presence of Mg^2+^ (Extended Data Figure 6; Extended Data Table 1). The structure harbours a strong resemblance to its ATP-bound counterpart, including the engagement of DNA binding loops with the tracking strand (Extended Data Figure 7a-c). However, moderate rotational differences in the C-tiers and their respective DNA binding loops are observed – most notably in the F subunit, which is inherently more flexible. As a result, the DNA is slightly shifted compared to the ATP-bound structure (Extended Data Figure 7a). Every subunit is associated with an AMP-PNP molecule, its γ-phosphate coordinating with the Mg^2+^ cofactor (Extended Data Figure 7b). The density of AMP-PNP at the F subunit appears slightly diminished (Extended Data Figure 7a). This could be attributed either to inherent flexibility of the entire subunit, or reduced nucleotide occupancy. Analysis of the interfaces classifies A/B, B/C and C/D as ATP-like, and D/E and E/F as ADP-like (Extended Data Figure 7b). The interfaces generally seem tighter relative to the ATP-bound structure, in particular the C/D interface which exhibits an altered nucleotide coordination (Extended Data Figure 7b). Cumulatively, when examining LTag assemblies whereby hydrolysis is prevented, the data suggest that ATP hydrolysis takes place amid subunits toward the top of the staircase, while ADP release occurs at those near its base.

### Continuous reconstruction of a AAA+ helicase in action

To gain insight into the dynamic mechanism of hydrolysis-coupled DNA translocation and unwinding by a helicase, we reconstituted LTag with a fork substrate and ATP, and initiated hydrolysis by adding Mg^2+^. This mixture was vitrified and analyzed using cryo-EM. Image processing yielded a consensus reconstruction at 3 Å resolution (Extended Data Figure 8; Extended Data Table 2-4). When compared to non-hydrolytic structures, an overall increase in disorder is evident (Extended Data Figure 8), being particularly pronounced in specific regions like the DNA as well as subunits A and F. This is indicative of the reconstruction embodying an averaged superposition of different conformational states arising from ATP hydrolysis. Density corresponding to the fork nexus is discernible yet weak, likely due to the averaging effect of various hexamer positions around the nexus, combined with instances of LTag translocating along DNA strands from fully unwound substrates. To separate these averaged states, the dataset was subjected to 3D variability analysis (3DVA) ^28^ with a specification of three principal components (Extended Data Figure 8). Five volumes spanning approximately equally spaced points along each of the principal components were robustly modelled by means of interactive molecular dynamics flexible fitting ^30^ after further post-processing with EMReady ^29^ (Extended Data Table 2-4). Morphing along these volume trajectories illustrates the dynamic actions of LTag as it translocates on DNA (Supplementary Videos 1-3). Global atom resolvability scores (Q-scores ^31^) of the refined models approach average values reported for models built into high-resolution (∼3 Å) cryo-EM maps ^31^ (Extended Data Table 5), enabling the detailed tracking of molecular interaction changes throughout the process. Furthermore, geometric criteria for all the models lie in the top (0^th^-1^st^) percentile of PDB structures, with map-to-model correlation scores in the region of 90%.

### Direct visualization of ATP hydrolysis coupled to DNA translocation

In the reaction landscape captured by the first variability component, the N-tier collar of the helicase remains largely fixed, providing a stable base, while the C-tiers engage in mutually coupled rotational movements (Figure 3a). These rotations are tightly correlated with the longitudinal movement of DNA, providing an explanatory mechanism for guided linear traversal of DNA through the helicase (Figure 3b-d). The C-tiers, along with their DNA binding loops, rotate in a unified manner, acting as single, rigid units (Figure 3d). In the initial phase of the trajectory, we observe concerted interactions between the DNA and helicase subunits A through E (Figure 3b-c). Specifically, residues H513 and K512 engage with their respective nucleobases and phosphate groups (Figures 3b-c). Progressing along the trajectory, there is a notable shift in interactions; the loop from subunit A disengages entirely from the top DNA base, while the loop of subunit B exhibits a reduced interaction with its nucleobase (Figure 3c). This is also accompanied by an increase in B-factors of the said loops and bases. In tandem, subunit F loop forms a stronger connection with the nucleotide at the bottom of the staircase (Figure 3c), accompanied by increased resolvability of the loop and the bottom base. This sequence of coordinated events collectively propels the DNA forward by one base through the LTag channel per hydrolysis event (Supplementary Video 4), as explained below.

**Figure 3.**
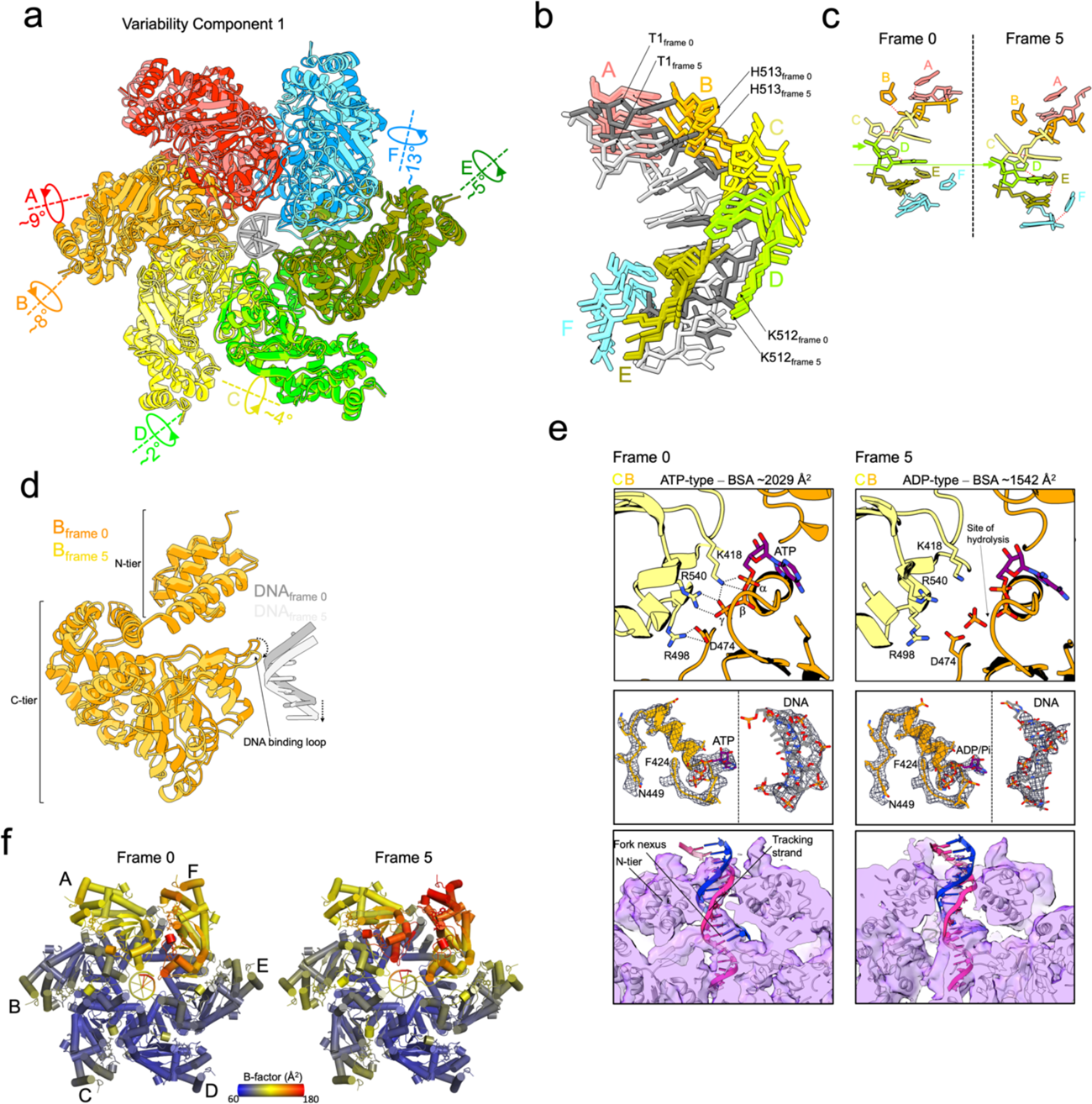
Visualization of ATP hydrolysis-coupled DNA translocation. **a** Model overlay of the initial and final frames along the first variability component, illustrating the extent and orientation of C-tier rotations in each subunit. **b** Overlay of DNA models from the initial and final frames of the first component, alongside the superimposed models of DNA-coordinating residues K512 and H513 across the five frames. **c** The initial and final frame models of DNA and the H513 side chain from the first component displayed side by side. Nucleotides are color-coded to match the associated DNA binding loop. Hydrogen bonds between H513 and nucleobases are depicted with dotted red lines. Green arrows indicate the phosphate group of the nucleotide engaged with the DNA binding loop of subunit D, emphasizing the net translocation of one base along the trajectory. **d** B subunit models from the initial and final frames of the first principal trajectory superimposed to show C-tier rotation, which causes a downward shift of both the DNA binding loop and the DNA it binds. **e** *Top insets*: Interactions at the nucleotide binding pocket between the B and C subunits in the initial and final frames of the first component. Calculated Buried Surface Areas (BSA) are shown, along with the assigned interface type. Hydrogen bonds are shown as black dotted lines. The analysis reveals a disruption at the interface between subunits B and C, indicative of a hydrolytic event. The location where ATP hydrolysis is proposed to occur is highlighted. *Middle insets*: Focus on map density around the nucleotide binding pocket and DNA regions in the first and final frames of the first component, demonstrating the 3DVA maps’ quality. *Bottom insets*: Unfiltered cryo-EM map and model of the first and final frames of the first component. The map is displayed at a low threshold to reveal the density throughout the N-tier channel, suggesting the presence of double-stranded DNA. **f** Models of LTag from the initial and final frames of the first component colored according to B-factors, highlighting an increase in overall B-factors in the final frame, especially within the A, B, and F subunits.

The analysis of inter-subunit interface evolution provides insight into the link between ATP hydrolysis and DNA translocation (Figure 3e). Across the trajectory, the B/C interface – which starts as the most stable – shows dramatic changes, with a ∼25% reduction in interface area and a ∼40% decrease in contact points by the end (Figure 3e). Closer examination reveals that this interface transitions from an ATP-bound state to an ADP-bound state, marked by the complete disassociation of the finger R540 from the γ-phosphate of ATP (Figure 3e). The A/B interface, in contrast, shows only a minor decrease in its buried surface area (∼9%) and reduction of only one contact point (Extended Data Figure 9a). This slight change leads to a moderate disturbance of the interface, where its ATP-type conformation becomes partially altered due to an increased gap between R540 and the γ-phosphate, along with a disruption of the R498-D474 salt bridge (Extended Data Figure 9a). The other interfaces show minimal changes in nucleotide coordination, but nonetheless undergo rotations in response to motions in upstream subunits (Figure 3a; Extended Data Figure 9a). The divergence in dynamics between the A/B and B/C interfaces correlates with the rotational alignment of the subunit C-tiers: A and B rotate in concert, while B and C rotate asynchronously, leading to full disconnection at the B/C interface (Figure 3a). Throughout the trajectory at the B/C interface, cryo-EM density consistent with the γ-phosphate of ATP is present despite changes in the coordination of interacting residues (Extended Data Figure 9a). This may be either due to replacement of the γ-phosphate by a chloride ion after ATP hydrolysis, as seen in the ADP-bound LTag structure (Extended Data Figure 5), or retention of the hydrolyzed phosphate within the binding pocket through interactions with both cis- and trans residues. MD simulations suggest the hydrolyzed phosphate group remains trapped in the nucleotide pocket (Supplementary Video 5). Notably, a recent investigation combining cryo-EM, NMR and MD simulations identified a metastable ADP·Pi reaction intermediate in the AAA+ enzyme p97 ^32^, supporting the hydrogen phosphate group remaining confined inside the pocket post-hydrolysis. ADP+Pi would then eventually be released when it reaches the bottom of the staircase, with its far fewer supporting interactions. Importantly, when the cryo-EM maps are examined at a lower threshold, features corresponding to double-stranded DNA become apparent at the fork nexus (Figure 3e). This observed density pattern supports that the dynamic movements captured represent DNA unwinding occurring within the inner chamber of the helicase.

The first principal component highlights ATP hydrolysis at the B/C interface as a crucial molecular event, functioning like an ’entropy switch’. Disengagement of the catalytic finger Arg540 from ATP γ-phosphate triggers a transition from a more ordered, strained state, to a less ordered, more relaxed state within the subunits. The transition, which is characterized by a discernible increase in overall atomic B-factors particularly in subunits A and B (Figure 3f), reflects the entropic release and consequent onset of C-tier rotations relative to the N-terminal collar domains, underpinning DNA strand progression. Significantly, these movements take place without direct allosteric changes to the DNA binding loops, which remain attached to the DNA throughout. The analysis thereby implies that energy from ATP hydrolysis does not directly drive translocation, but rather removes a block to translocation. Following this, the second principal component illustrates the preparatory steps of reconfiguring the loop staircase, priming the system for the subsequent cycle of ATP hydrolysis and translocation.

### Dynamics of preparatory steps for translocation cycle restart

Similar to the first component, the second principal trajectory marks a pre-hydrolysis phase at the outset (Figure 4). Progression through this trajectory reveals three key events: firstly, subunit A release from DNA; secondly, DNA advancement through the channel; and thirdly, detachment of subunit F’s loop from DNA and its subsequent movement upward along the staircase, occurring in tandem with closing distance between subunits F and A (Figure 4a-c; Supplementary Video 6). The rotational axes of the C-tiers exhibit variations from those observed in the first component (Figures 3a and 4a). Notably, the axes for subunits A and B demonstrate reduced collinearity, and the F subunit rotation is in the opposite direction – towards subunit A (Figure 4a). Interfacial analysis indicates disruptions at both A/B and B/C interfaces. The observation might suggest dual hydrolytic events; however, we deduce that A/B interface disruption is not a primary hydrolysis event but rather a consequence of hydrolysis at the B/C interface and resulting rotation of the B subunit C-tier away from that of A. This argument is supported by the intrinsic propensity of the A/B interface for motion, due to subunit A’s limited stabilizing contacts with the previous F subunit, which is also reflected in the elevated B-factors for the C-tier of the subunit A compared to subunit B (i.e. 114 versus 88 Å^2^). Furthermore, unlike subunit C, the subunit B’s ATP-coordinating residue R498 exhibits no side chain density in the trajectory’s initial frame, and is not engaged in a stable salt bridge with D474 (Extended Data Figure 9b). In the crystal structure of LTag bound to ATP without DNA ^10^, R498 is pivotal for positioning the “apical” water molecule critical for hydrolytic activity at the active site. This water molecule, situated at the metal ion coordination sphere apex, facilitates the nucleophilic attack on ATP γ-phosphate, thereby catalyzing its hydrolysis. The increased P-value for the A/B interface, when compared with the B/C interface (Figure 3d), further supports its inherently dynamic nature and less supportive state for ATP hydrolysis. Hence, it is highly likely that the A subunit, while releasing from B, is still ATP-bound. As the cycle progresses, the former A/B interface, now in the B/C position, will achieve the necessary stability to undergo ATP hydrolysis in a subsequent translocation cycle.

**Figure 4.**
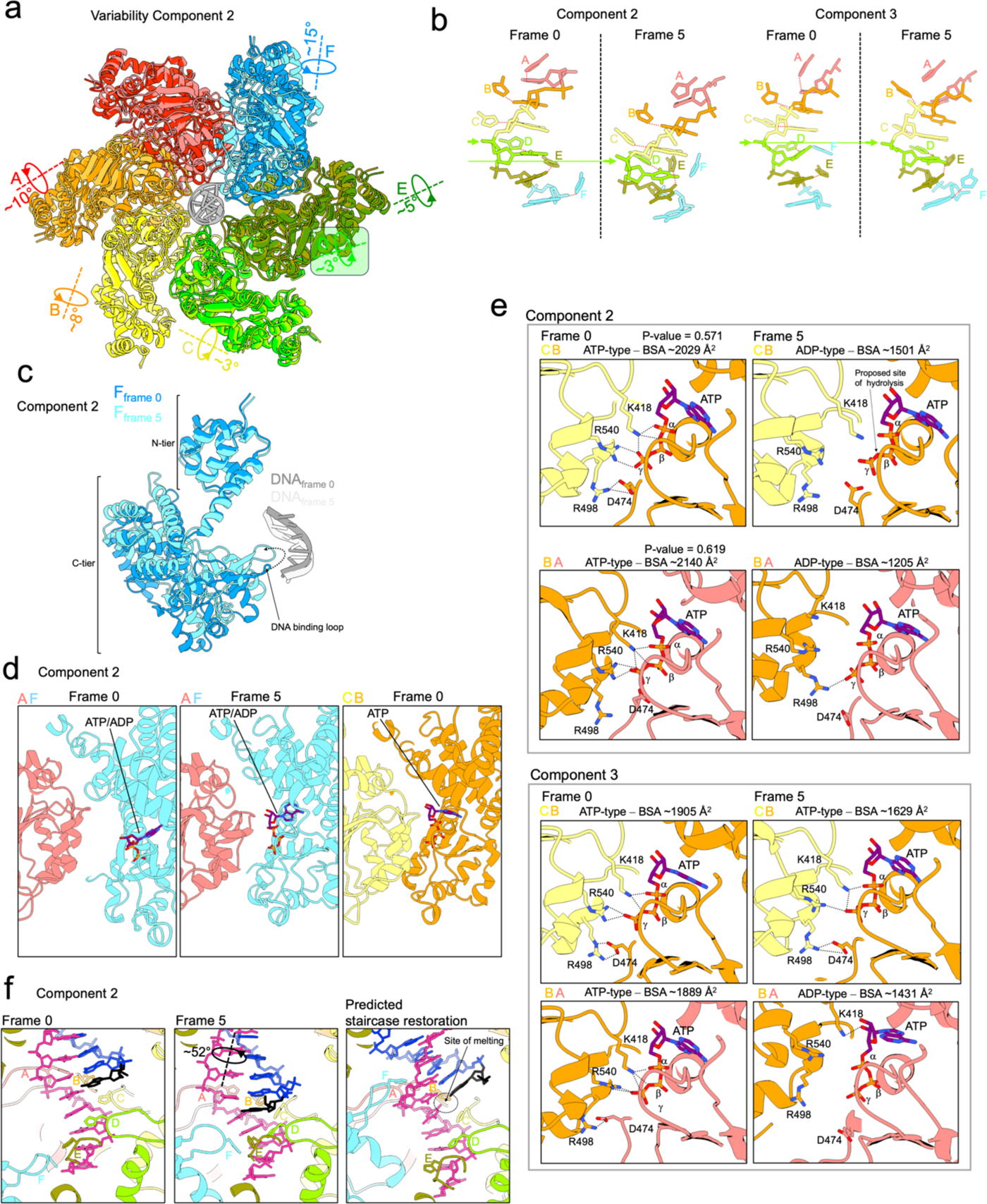
Dynamics of preparatory steps for hydrolysis and translocation cycle restart. **a** Overlay of models from the initial and final frames of the trajectory, as derived from the second variability component, illustrating the magnitude and direction of C-tier rotations in each subunit. **b** Initial and final frame models of DNA and the H513 side chain from the second and third components are displayed side by side. Hydrogen bonds between H513 and the nucleobases are depicted with dotted red lines. Green arrows indicate the phosphate group of the nucleotide engaged with the DNA binding loop of subunit D, showing the net translocation of one base along the second principal component, and no translocation along the third component. **c** Overlay of F subunit models from the initial and final frames of the second principal component traversal, showing the C-tier rotation and upward shift of the DNA binding loop, which results in the detachment of the DNA from the subunit. **d** Interfaces between the C-tiers of the A and F subunits in the initial and final frames of the second component. A third inset presents the interface between the C-tiers of the B and C subunits in the initial frame, emphasizing the narrowing gap between the A and F subunits throughout the trajectory. **e** Interactions at the nucleotide binding pocket between the B and C subunits, or A and B subunits in the initial and final frames of the second and third variability components. Calculated Buried Surface Areas (BSA) or P-values are shown, alongside the assigned interface type. Hydrogen bonds are shown as black dotted lines. The analysis reveals a disruption at both BC and AB interfaces in the second component, and no disruption of the BC interface in the third component. These findings collectively indicate that disruption of the BC interface is necessary for DNA translocation. **f** Extrapolation of the conformational changes at the DNA fork nexus in a full translocation cycle. The dsDNA segment is modelled based on the pre-hydrolytic structure (Figure 1). The structure in the third panel is equivalent to the initial frame of the second component, but with the LTag subunits rotated one step around the six-fold symmetry axis of the N-tier. The DNA nexus rotation required to re-establish the loops staircase leads to the unpairing of the terminal base of the passive strand (colored in black).

Disengagement of the DNA binding loop of subunit F from DNA and its movement up the staircase, along with the narrowing gap between the F and A subunits, signifies the preparatory phase for the next cycle of translocation. The very last steps, e.g. F subunit’s de facto capture of a new nucleobase likely occur too quickly to be resolved by 3DVA, with the end result indistinguishable to the starting state.

Analysis of the third component (Supplementary Video 7) reinforces the notion that the B/C interface breakage is essential for DNA translocation (Figure 4b and 4e). Here, the DNA remains static (Figure 4b), with the B/C interface retaining its ATP-bound conformation. Meanwhile, subunit F appears to be bound to ATP at either end of the trajectory. However, A establishes a stable interaction with DNA (Figure 4b), and via ATP with B, when approached by an ascending ATP bound subunit F (Figure 4e).

Integrating insights from these analyses with that of the helicase in its pre-hydrolysis state (Figure 1), allows us to delineate the series of conformational changes occurring at the DNA fork nexus throughout a complete ATP hydrolysis cycle (Figure 4f, Supplementary Video 8). During this cycle, the nexus moves within the helicase inner chamber, driven by the motion of the DNA binding loops that pull on the tracking strand. Concurrently, as subunit F and its loop moves upward, restoring the staircase structure, the nexus rotates. This rotation is triggered by the incorporation of the free nucleobase at the top of the staircase into the compact helix of the tracking strand. The section of the tracking strand gripped by the loops instead translates without rotation. This differential movement leads to the melting of the base pair at the fork junction (Figure 4f). Here, the terminal base of the passive strand is unable to engage with the counterpart on the tracking strand, which is coordinated by H513 of the B subunit.

### Generalization of the entropy-switch mechanism for substrate translocation

The mechanistic basis of hydrolysis-coupled translocation by LTag helicase is likely to be generally applicable to other hexameric helicases believed to operate via a rotary mechanism, irrespective of different step sizes or polarity. These include the viral E1 helicase ^25^, bacteriophage T7 helicase ^27^, bacterial DnaB ^21^ and Rho ^23^, as well as archaeal MCM ^24^. By contrast, the heterohexameric structure and functional asymmetry^7^ of eukaryotic CMG have inspired models of asymmetrical translocation ^26,33^. Initially, only three ssDNA-bound CMG conformers were identified, fewer than the six expected for symmetric translocation ^26,34^. However, recent cryo-EM studies have revealed two additional conformers ^35^. Thus, considering the conserved ssDNA-binding residues across all MCM hexamers ^24^, it is likely that CMG translocates on DNA via a universal, symmetric rotary mechanism. Furthermore, our interpretation of ATP hydrolysis as an entropy switch could explain why disabling specific ATPase subunits in CMG does not drastically affect DNA translocation/unwinding capability ^7^. For example, a Drosophila CMG mutant with a KA alteration in the Walker A motif of MCM6 retains approximately 70-80% of the unwinding efficiency of the wild type, whereas the same mutation in MCM3 reduces the activity to about 1-5% ^7^. In the structure of human CMG bound to forked DNA, MCM6-4 and MCM3-5 are the strongest and weakest interfaces, respectively ^22^. Notably, in a yeast CMG conformer with the MCM6 subunit at the top of the staircase ^35^, MCM6 lacks an associated nucleotide, indicating the inherent stability of the MCM6-4 interface even without ATP. Our model suggests that ATP binding rebuilds the staircase, and hydrolysis releases the translocation brake. Thus, a mutated MCM6 might ascend the staircase and re-establish the MCM6-4 interface without ATP due to its intrinsic affinity. Following one translocation cycle, the mutant MCM6-4 interface would be primed for hydrolysis, but translocation proceeds as the “block” is already absent. Interestingly, AlphaFold3 ^36^ predictions show KA mutations in Walker A significantly affect ATP-binding by MCM3, but not by MCM6; which may further help explain mutation tolerance by the latter. Furthermore, we would like to note that operating principles of the entropy-switch mechanism may indeed be relevant to other hexameric AAA+ substrate translocases, e.g. VAT or ClpXP ^37,38^.

### A general mechanism for DNA unwinding

The structure of LTag bound to forked DNA and ATP, in the absence of the Mg^2+^ ion, captures a critical stage in the unwinding process, positioning the fork nexus within the helicase domain. The wedge created by the DNA binding loops and the restricted inner diameter of the helicase obstructs duplex DNA from completely traversing the C-tiers intact, thereby serving as the central site for DNA melting. Critically, the overlay of the B and C subunits from this structure onto the crystal structure of the LTag dimer bound to the early palindrome (EP)-half of origin DNA ^9^, demonstrates a striking alignment in the region across the fork nexus and DNA binding loops (Figure 5a). Such a comparison suggests that the initiation of origin melting is contingent upon the sequential assembly of the remaining LTag subunits and ATP binding (Figure 5a). This assembly would complete the helicase loop staircase, with the D, E, and F subunits loops penetrating the DNA minor groove to encircle the tracking strand fully (Figure 5a, inset). Notably, the assembly of LTag with origin DNA in the presence of ATP has been shown to initiate melting at the EP sequence, spanning 8 base pairs ^39^. These bases align with the origin DNA segment that would be laterally displaced by the loops of the D, E, and F subunits in the fully assembled LTag hexamer, providing a mechanistic correlation to the melting process (Figure 5a). It is probable that a comparable melting event happens at the AT-rich half of the origin. Here, the formation of a second hexamer was shown to cause distortions in the AT-rich area, located symmetrically from the origin center, similar to the EP-region ^39^. Consequently, after melting these areas, each LTag hexamer would be in a configuration ready for ATP-driven translocation.

**Figure 5.**
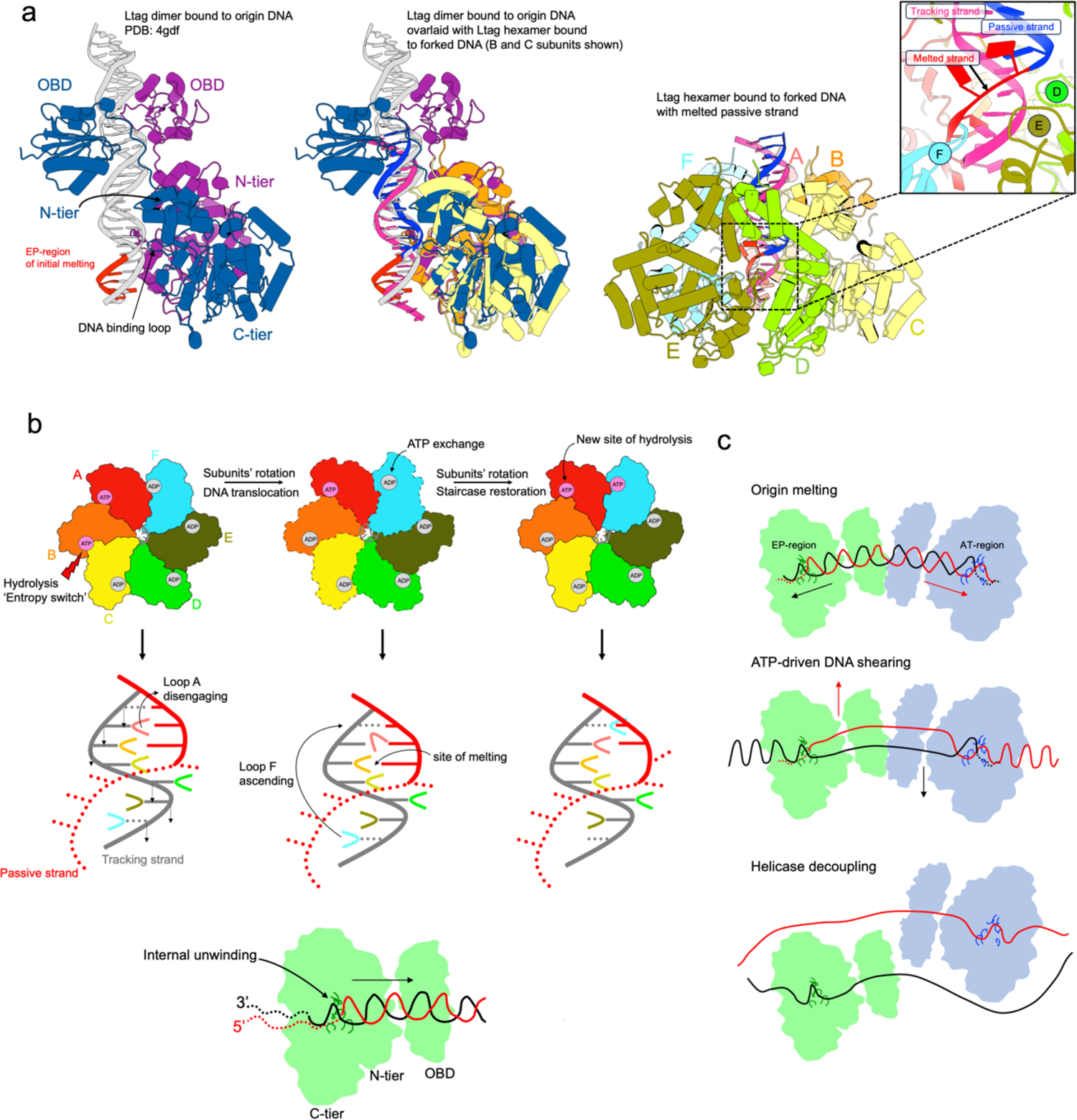
Model for origin melting and DNA unwinding. **a** *Left panel*: Crystal structure of Ltag dimer bound to origin DNA ^9^. The early-palindrome (EP) region of origin DNA, which undergoes initial melting upon LTag double hexamer assembly ^39^ is colored red. *Middle panel*: Superposition of the structure showed in the left panel and the cryo-EM structure of LTag bound to forked DNA determined in this work. Only the B and C subunits of the latter structure are shown. *Right panel*: The cryo-EM structure of LTag bound to forked DNA, with the passive strand extended by 6 bases colored red, paralleling the DNA origin-melting process; demonstrating how the intrusion of loops from D, E, and F subunits into the DNA minor groove – to fully engage the tracking strand – leads to the unpairing of DNA bases. **b** Mechanism of internal DNA unwinding by LTag, synthesizing insights from pre-hydrolysis structure of LTag bound to forked DNA and ATP, with continuous heterogeneity analysis. The model outlines a single translocation cycle whereby ATP hydrolysis at the hexamer tightest interface (BC) acts as an ’entropy switch’. Hydrolysis initiates synchronized rotations of the C-tiers across different subunits and rearrangements of the DNA binding loops, facilitating longitudinal movement of the tracking strand and DNA nexus within the LTag inner chamber. Following the exchange from ADP to ATP by the F subunit, a new series of C-tier rotations occurs. This shifts the DNA binding loop of the F subunit to the top of the staircase, concurrently establishing the ATP interface between the F and A subunits. The reconstitution of the staircase structure leads to the melting of one base pair at the nexus, progressing the unwinding process. **c** Model of origin unwinding facilitated by dual LTag helicases: assembly of two hexamers in a head-to-head configuration at the origin causes melting at the EP and AT-rich regions. The ATP-driven traction on the tracking strand by each hexamer leads to shearing of the DNA between the hexamers and the creation of single-stranded loops at the rear of the helicase. When the unwound DNA strands spanning the hexamers are fully elongated, they exit their respective hexamers. This results in the passive strands being routed around the exterior of the hexamers while concurrently releasing the tracking strand loops formed at the back of the helicase, culminating in separation of the helicases.

Our data suggest that translocation and unwinding can occur within the helicase domain (Figure 4f, Figure 5b and Supplementary Video 8). This model is in strong agreement with previous biochemical evidence that LTag can unwind forked substrates lacking a 5’ tail ^40^. Critically, it was determined that a minimum of 5 nucleotides in the 3’ tail is necessary for efficient unwinding ^40^, matching the number of nucleotides needed to engage all the loops in the staircase (Figure 1). However, internal unwinding might not always be the most efficient process: as LTag advances, the melted passive strand, exiting from the back of the C-tiers, could potentially re-anneal with the tracking strand ^18^. Nevertheless, when two opposing hexamers collide at the origin, they each would pull their respective tracking strand in opposite directions ^18^ (Figure 5c). This action, over multiple hydrolysis events, would shear and unwind the strands between the hexamers while concurrently twisting them beyond the DNA-binding loops, effectively preventing their re-annealing and leading to the formation of single-stranded loops extending from the C-tiers (Figure 5c). Once the unwound DNA strands between the hexamers are fully stretched, they would be expelled from their corresponding hexamers, resulting in passive strands running outside the hexamers and the simultaneous release of the tracking strand loops formed at the back of the helicase (Figure 5c). Passive strand lateral pulling and ejection are likely facilitated by the curvature observed in the longitudinal axis of the LTag double hexamer bound to origin DNA captured by EM ^13^. At this stage, the two uncoupled hexamers would be poised for DNA unwinding through steric exclusion ^19,20^. The passive strand may be ejected through the large gap between the C-tiers of the top and bottom subunits of the staircase. However, transient gaps between interfaces in both the N-tier collar and OBD ring are also necessary for this process. The known ability of LTag to oligomerize from dimers to hexamers ^15,16^ supports the feasibility of these gap formations. In eukaryotic systems, as in the SV40 system, CMG helicases arranged head-to-head at origins lead to the formation of bidirectional replication forks ^6,41^. The structure of yeast CMG double hexamer bound to duplex DNA ^42^ and that pertaining to the initial phase of origin melting ^43^ indicate that origin melting is fundamentally driven by ATP binding, DNA binding loop staircase formation, and tracking strand engagement by the loops (Extended Data Figure 10). These factors, as proposed for the SV40 system, contribute to strand separation within the helicase inner chamber and enable each hexamer to acquire a configuration primed for ATP-driven translocation ^44^. As such, principles underlying origin melting appear conserved in both domains (Extended Data Figure 10). Considering CMG likely translocates on DNA via a general rotary mechanism ^24,35^, it is plausible that DNA shearing and helicase uncoupling proposed here remains applicable.

## Methods

### Protein cloning, expression, and purification

Full-length SV40 large T antigen (LTag) (accession number: P03070) N-terminally double 6X histidine-tagged along with TEV cleavage site sequence was cloned into pFastBac1 plasmid by GenScript. The recombinant expression plasmid was introduced into MultiBac baculoviral DNA in DH10MultiBac™ and bacmid DNA was isolated. To prepare the baculovirus, the bacmid DNA of Ltag was transfected into Sf9 cells using FuGENE® HD (Promega) according to the manufacturer’s instructions. The resulting supernatant was obtained as the P1 virus stock which was then amplified to obtain P2 virus stock. P2 virus stock was then further amplified to obtain P3 virus stock for large-scale expression. LTag was expressed by transfecting 4 L of Sf9 suspension culture at a density of 2 × 106 cells/mL with LTag P3 virus for 55-60 hrs. Cells were harvested by centrifugation at 5500×g for 10 min and re-suspended in lysis buffer [20 mM Tris (pH 8.0), 50 mM Imidazole, 500 mM KCl, 1 mM Dithiothreitol, 5% (v/v) Glycerol and EDTA-free protease inhibitor cocktail tablet/50 ml (Roche, UK)]. The cell lysate was clarified by centrifugation at 95,834×g for 1 hr at 4 °C. The supernatant was directly loaded onto a 5 ml Histrap affinity column (Cytiva) pre-equilibrated with buffer A [20 mM Tris (pH 8.0), 50 mM Imidazole, 500 mM KCl, 1 mM DTT and 10% Glycerol]. The loaded column was washed with 50 ml of buffer A containing 50 mM imidazole followed by 50 ml of Buffer A containing 100 mM imidazole to remove the non-specific binding of the protein to the column. Finally, the protein was eluted with a 20 ml gradient to 500 mM imidazole using Buffer B containing low salt [20mM Tris (pH8.0), 500 mM Imidazole, 250 mM KCl, 1mM DTT, 10 % Glycerol]. The pooled fractions of LTag were treated with TEV protease to remove the His-tag and dialyzed overnight in dialysis buffer [20 mM Tris (pH 8.0), 100 mM KCl, 1 mM DTT, and 10% glycerol]. The cleaved protein was then loaded into Histrap HP 1 ml affinity column (Cytiva) pre-equilibrated with Buffer C [20 mM Tris (pH 8), 250 mM KCl, 1 mM DTT and 10% Glycerol]. The loaded column was then washed with 10 ml of Buffer D followed by a 10 ml linear gradient of buffer B. Flow-through fractions that contain LTag were concentrated and loaded onto a HiLoad 16/600 Superdex 200 pg (Cytiva) pre-equilibrated with gel filtration buffer [20 mM Tris (pH 8.0), 100 mM KCl, 1 mM DTT and 10% Glycerol]. Collected fractions were flash-frozen and stored at −80 °C.

### DNA substrates

The forked DNA substrate for reconstitution with LTag and ATP in absence of Mg^2+^ was generated by annealing the oligonucleotides 5’-GGTGATCGTTCGCTACATGTCGTCAGGATTCCAGGCAG-3’ and 5’-AGACACATGGATGTAGCGAACGATCACC-3’ purchased from Eurofins. The forked DNA substrate for reconstitution with LTag and ADP, LTag and ATP in the presence of Mg^2+^, or LTag and AMP/PNP in the presence of Mg^2+^ was adapted from SenGupta & Borowiec ^45^, with unpaired arms changed to poly-Ts. The constituent oligonucleotides were therefore 5’-TTCTGTGACTACCTGGACGACCGGGTTTTTTTTTTTTTTTTTTTTTTTTTTTTTT-3’ and 5’-TTTTTTTTTTTTTTTTTTTTTTTTTTTTTTCCCGGTCGTCCAGGTAGTCACAGA-3’.

The annealed product was purchased from Eurogentec. The gap DNA substrate for reconstitution of LTAg and ATP apo complex was created by annealing oligonucleotides 5’-AGCTATGACCATGATTACGAATTG[23ddC]-3’ and 5’-TTTTTCGGAGTCGTTTCGACTCCGATTTTTTTTTTTTTTTTTTTTTTTTTTTTTTTTTTTTTTTTTTTTTGCAATTCGTAATCATGGTCATAGCT-3’. LTag showed no binding to this substrate without a free 3’ ssDNA tail.

### Buffers

The buffer used for LTag-ATP-DNA and LTag-ADP-DNA and LTag-ATP (apo) samples was based on DNA binding assay from Li *et al.* ^11^ comprising 1 mM ATP, 50 mM Tris-HCl (pH 8.0) and 200 mM NaCl. The buffer used in the LTag-ATP-Mg^2+^-DNA sample was based on DNA helicase assay from ^11^, composed of 50 mM HEPES (pH 7.5), 1 mM ATP, 3 mM MgCl_2_, 1 mM DTT and 50 mM NaCl. The buffer used for the LTag-AMPPNP-Mg^2+^-DNA complex was 50 mM Tris-HCl, pH 7.5, 1 mM DTT, 100 mM NaCl, 5 mM MgCl and 3 mM AMP-PMP.

### Cryo-EM sample preparation

The LTag stock was pre-incubated with buffer before addition of DNA substrate (in equimolar ratio to putative hexamer). The steps were performed in a 4 °C temperature-controlled room. The final protein concentration in samples applied to the grids was ∼0.098 mg/ml (equivalent to 0.2 µM hexamer). R1.2/1.3 or R2/2 UltrAuFoil^®^ grids (Quantifoil Micro Tools GmbH) were glow-discharged with 40 mA for 5 min inside a Quorum GloQube^®^ Plus unit. Graphene oxide (Merck KGaA) coating was applied^46^. 3 µl of sample was applied to the grid in a VitroBot Mark IV at 4 °C and 100% relative humidity, and following a wait time of 30 s, blotted for 3s with blot-force 4, and plunged into liquid ethane. For the LTag-AMPPNP-Mg^2+^-DNA complex, 2 μM LTag (with respect to the hexamer) was mixed in the corresponding buffer and incubated at 4 °C for 30 min. Following this, 2 μM DNA was added and the reaction incubated at 4 °C for a further 30 min. 3 uL of sample was deposited onto a R2/2 Au 300 mesh grid previously glow discharged at 40 mA for 60 sec. The grid was frozen on a Vitrobot with a 3 sec blotting time, 0 sec wait time and a blot force of 5, with 4 degrees C and 100% humidity in the chamber.

### Cryo-EM data acquisition

Data for the LTag-AMPPNP-Mg^2+^-DNA structure were collected on a Titan Krios G4 TEM (Thermo Fisher Scientific, Eindhoven) located at the Imaging and Characterization Core Lab of the King Abdullah University of Science and Technology, Saudi Arabia. Movies in EER format were recorded at 130,000x magnification (calibrated pixel size of 0.93 Å px^-1^) using a Falcon 4i direct detection camera at the end of a Selectris-X energy filter with a 10eV slit width. Data were collected with a defocus range of -2.5 to -1.5 μm and an accumulated dose of 38.7 e-/Å^2^ . All other data were collected on a Titan Krios G3 TEM (Thermo Fisher Scientific, Eindhoven) at 105,000× magnification in AFIS mode with 100 mm objective aperture, using the GATAN K3 direct detection camera at the end of a GATAN BioQuantum energy filter (20 eV slit width) in counted super-resolution mode (binning 2) at the Leicester Institute of Structural and Chemical Biology, UK. Individual exposures were acquired at a dose rate of 18 e^−^ px^-1^ s^-1^ in 50 equal fractions over 2 s, with a calibrated pixel size of 0.835 Å px^-1^ and an accumulated dose of 50 e^−^ Å^-2^. The nominal defocus was varied between -1.0 µm and -2.5 µm in 0.3 µm steps. The gross number of movies collected from each sample were: 8,072 (LTag-ATP-DNA); 9,737 (LTag-ADP-DNA); 6,127 (LTag-AMPPNP/Mg^2+^-DNA); and 8,460 (LTag-ATP/Mg^2+^-DNA).

### Cryo-EM Data processing

Movie stacks were corrected for beam-induced motion using RELION’s implementation^47^ of MotionCor2^48^ with dose-weighting, and CTF estimation of the integrated micrographs was performed using CTFFIND 4.1^49^. The micrographs were denoised prior to picking by Topaz^50^, both using default models. The picks were extracted using a box size of 384, and subject to several iterations of 2D classification and selection using VDAM in RELION 4^47^ (with decreasing binning factors of 4×, 2× and 1×). In general, particle subsets from above were imported into cryoSPARC (Structura Biotechnology Inc., Toronto, Canada) and re-extracted from Patch Motion Correction and Patch CTF Estimation jobs, and initially subjected to *ab-initio* reconstruction with 3 classes (allowing for DNA-bound, apo and junk categories). Where two junk classes were detected, the process was repeated using 2 classes. The resulting particle selections were further corrected for beam-induced motion with Local Motion Correction. Eventual final reconstructions were derived using non-uniform refinement, with CTF and per-particle scale refinements enabled. The LTag-ATP-DNA reconstruction was initially refined to 2.9 Å in RELION 4 using SIDESPLITTER. The final particle subset following 2D classification was subject to a 3-class 3D (VDAM) initial model job. This produced two junk classes; the non-junk class was subjected to 2-class 3D classification. This yielded apo (54%) and DNA-bound (44%) classes. The latter was iteratively 3D auto-refined employing CTF refinements and Bayesian polishing. The refined particle stack with its assignments was imported into cryoSPARC, and re-refined using non-uniform refinement. This gave a better regularised map despite a slightly lower reported global GSFSC resolution. 3D variability analysis^28^ of the LTag-ATP/Mg^2+^-DNA consensus reconstruction was carried out using its refinement mask (created as soft mask around protein and bound DNA); with 3 modes and filter resolution of 5 Å (the global GSFSC resolution being 3 Å), as well as a high-pass resolution of 150 Å. Latent coordinates exhibited a spherical/3D Gaussian distribution, characteristic of continuous conformational variability. 20 volumes (frame_000 to frame_019) were sampled along each principal component (0, 1, 2). Data had been exhaustively cleaned via repeat classification without separating conformations. The LTag-AMPPNP-Mg^2+^-DNA structure was wholly refined in RELION 4. Briefly, particles picked by Topaz (2,859,203 particles) were extracted with a box size of 256 pixels and 4X downscaling (3.72 Å/px), and extensively cleaned by 2D classification. The resulting particles (957,358 particles) were subjected to 3D classification with 3 classes. Particles from the best class (235,707 particles) were re-extracted to full size (0.93 Å/px) and subjected to 3D refinement, CTF-refinement and polishing, resulting in a 3.2 Å resolution map. LTag-ATP (apo) reconstruction from the gapped DNA sample was refined in RELION. Steps up to and including 2D classification and subset selection were done in RELION 4. Ab-initio reconstruction was obtained from cryoSPARC. 3D classification and auto-refinement was carried out in RELION 5 using Blush regularisation (with CTF refinement). Bayesian polishing was not performed. Total number of particles incorporated into the final reconstructions were: 92,330 (apo LTag-ATP); 210,910 (LTag-ATP-DNA); 97,587 (LTag-ADP-DNA); 235,707 (LTag-AMPPNP/Mg^2+^-DNA); and 201,416 (LTag-ATP/Mg^2+^-DNA). Local resolution maps were calculated in Phenix, and rendered on the full map (pre post-processing) with adjusted colouring. 3DFSC^51^ metrics were obtained from web server https://3dfsc.salk.edu/.

### Model building and refinement

The resulting volumes were post-processed with EMReady for purposes of interactive model building in ISOLDE^30^, alongside masked full/globally sharpened maps using automatically calculated B-factors. RELION or cryoSPARC post-processed maps were also used to derive draft models from ModelAngelo to guide the subsequent process. No form of post-processed maps were used for iterative map refinement itself. *Modelling of the LTag-ATP-DNA complex:* Initially, the crystal structure of ATP-bound LTag (1SVM) ^10^ was real space refined into the LTag-ATP-DNA map using PHENIX, and oligo-dT_6_ from E1-ADP-Mg^2+^ssDNA crystal structure (2GXA)^25^ fitted into density in ChimeraX. The map/model association was first globally simulated in ISOLDE, followed by multiple cycles of local rebuilding and Phenix real space refine with ISOLDE’s recommended settings. In latter rounds, additional bases in the tracking strand, and complementary strand were added and re-modelled. Several iterations of LocScale^52^ and ISOLDE refinement were used to improve the DNA nexus map and model. *Modelling of the LTag-ADP-DNA complex:* The crystal structure of ADP-bound LTag (1SVL)^10^ was flexibly fitted into the LTag-ADP-DNA reconstruction using iMODFIT and real space refined, with ssDNA incorporated as before. The model was then rebuilt and refined as previously. *Modelling of the LTag-AMPPNP-Mg^2+^-DNA complex:* A refined model of LTag-ATP-DNA (omitting the duplex portion of DNA) was initially rigid body fitted into the LTag-AMPPNP-Mg^2+^-DNA reconstruction in ChimeraX. Adaptive distance restraints were applied to individual chains in ISOLDE, and simulated against the map. Restraints were progressively released and model fully rebuilt. Towards the end, ATP was replaced with AMPPNP and Mg^2+^, and resimulated/rebuilt. *Modelling of the LTag-ATP/Mg^2+^-DNA complex into the 3DVA volumes:* The LTag-ATP-DNA model, following rigid body fitting into LTag-ATP/Mg^2+^-DNA consensus map, was first rebuilt into a working consensus model. This model was flexibly fitted into the central-most sampled volume (frame_010) from the first principal component. The top-ranked AlphaFold2^53^ monomer structure prediction for SV40 LTag was superimposed to each of the six subunits using MatchMaker^54^. Residues not pertaining to the core helicase 265-627 were deleted. Adaptive distance restraints were applied to individual chains, followed by a whole map-model simulation in ISOLDE. Restraints were progressively released, and the model rebuilt end-to-end. The derived model was placed into two intermediate volumes (frame_005 & frame_015). Adaptive restraints were reapplied and models entirely rebuilt as above. These models were placed into terminal volumes (frame_000 & frame_019), and rebuilt as per the same procedure. The central model from principal component 0 was placed into central volume of component 1. After remodelling, it was used as basis for modelling the two intermediate, and then terminal frames as before. The process was repeated for principal component 2. Each residue in all of the above models was manually inspected/rebuilt at least once.

Q-scores were calculated in Model-map Q-score^31^ ChimeraX plugin using default/recommended sigma = 0.6. The consensus LTag-ATP/Mg^2+^-DNA structure couldn’t be definitively modelled throughout due to representing multiple averaged states, and was used chiefly as an interim model to generate initial model drafts for variability volumes. Modelling of the LTag-ATP (apo) complex: LTAg-ATP (1SVM) crystal structure was first Phenix real-space-refined into the apo LTAg-ATP reconstruction postprocessed with EMReady. AlphaFold2 monomer prediction corresponding to amino acids 266 - 627 was superposed with each of the six chains using MatchMaker, and subjected to molecular dynamics bulk flexible fitting with adaptive distance restraints in ISOLDE. Restraints were released inside a whole map/model simulation, followed by localised rebuilding in ISOLDE. The final Phenix real space refined structures were assigned secondary structure annotations using DSSP ^55^.

### Map and model visualization

Maps were visualized in UCSF Chimera^56^ and ChimeraX^57^ and all model illustrations and morphs were prepared using ChimeraX or PyMOL.

### Statistics and reproducibility

No statistical methods were used to predetermine sample size. The experiments were not randomized, and investigators were not blinded to allocation during experiments and outcome assessment.

## Data availability

The data that support this study are available from the corresponding authors upon request. The maps of the LTag-ATP-DNA complex reconstituted without Mg^2+^ has been deposited in the EMBD with accession codes EMD-50289 and EMD-50287, and the atomic models in the Protein Data Bank under accession codes 9FB6 and 9FB4. The map of the LTag-ADP-DNA complex has been deposited in the EMBD with accession code EMD-50002, and the atomic model in the Protein Data Bank under accession code 9EVH. The map of the LTag-AMPPNP-DNA complex reconstituted with Mg^2+^ has been deposited in the EMBD with accession code EMD-50010, and the atomic model in the Protein Data Bank under accession code 9EVP. The consensus map of the LTag-ATP-DNA complex reconstituted with Mg^2+^ has been deposited in the EMBD with accession code EMD-50290. The 3DVA volumes have been deposited in the EMBD with accession codes EMD-50246, EMD-50245, EMD-50179, EMD-50190, EMD-50250 (Frames 1 to 5 of Component 1); EMD-50261, EMD-50257, EMD-50180, EMD-50256, EMD-50262 (Frames 1 to 5 of Component 2); EMD-50288, EMD-50265, EMD-50264, EMD-50244, EMD-50286 (Frames 1 to 5 of Component 3); and the atomic models in the Protein Data Bank under accession codes 9F75, 9F74, 9F3T, 9F5I, 9F7N (Frames 0 to 5 of Component 1); 9F9W, 9F9O, 9F3U, 9F9N, 9F9X (Frames 1 to 5 of Component 2); 9FB5, 9FA2, 9FA1, 9F73, 9FB0 (Frames 1 to 5 of Component 3).

## Contributions

T.S., S.M.H. and A.D.B conceived the study. M.T. and L.A. purified the protein. T.S., A.C., C.G.S. and A.D.B. performed cryo-EM imaging and processing. T.S. implemented the 3DVA analysis. T.S. and A.D.B. performed model building and validation. S.M.H. and A.D.B. supervised the study. T.S. and A.D.B. wrote the manuscript with contribution from the other authors.

## Acknowledgements

This research was supported by King Abdullah University of Science and Technology (KAUST) through core funding (to S.M.H. and A.D.B.) and the Competitive Research Award Grant CRG8 URF/1/4036-01-01 (to S.M.H. and A.D.B.). We thank past and present members of the S.M.H. and A.D.B. laboratories for help and discussions; Lingyun Zhao, Ashraf Al-Alamoudi, Alessandro Genovese and Rachid Sougrat for their assistance at the Imaging and Characterization Core Lab at KAUST; Tianyi Zhang for her help in revising the text. We acknowledge The Midlands Regional Cryo-EM Facility at the Leicester Institute of Structural and Chemical Biology (LISCB), major funding from MRC (MC_PC_17136). We thank T.J. Ragan (LISCB, University of Leicester) for his support with data processing and high-performance computing.

## Supplementary Information

### SI Guide

**Extended Data Figure 1.**
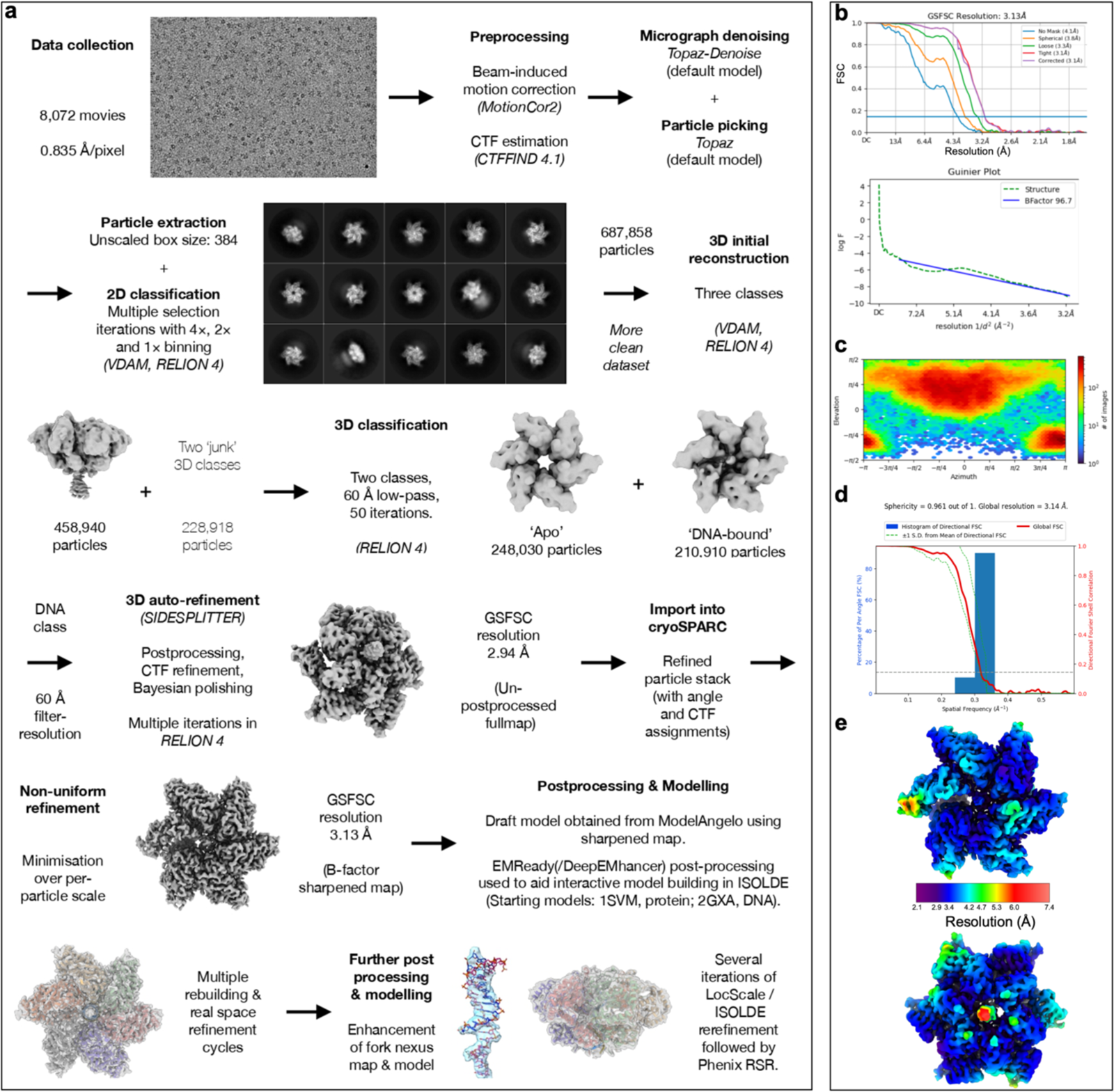
Cryo-EM of the LTag−ATP−DNA complex. **a,** Workflow schematics for cryo-EM data collection, processing, and modelling. Flows continue from left hand side of following row unless otherwise indicated **b,** Gold-standard Fourier shell correlation and Gunier plots **c,** Angular distribution of projections. Resolution is estimated using the 0.143 criterion. **d,** Map anisotropy analysis computed by 3DFSC^1^. **e,** Two views of the cryo-EM map colored by local resolution.

**Extended Data Figure 2.**
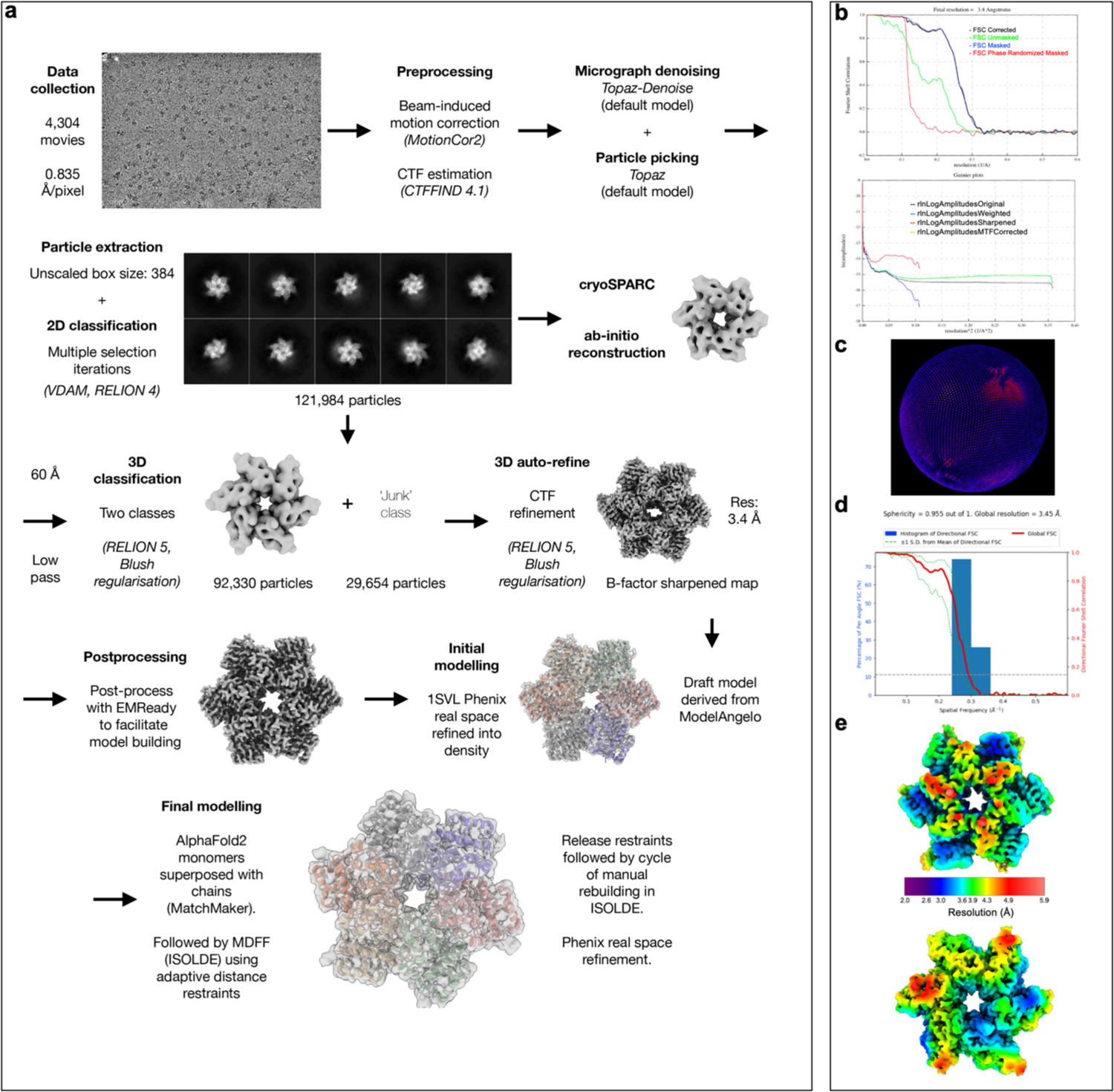
Cryo-EM of the LTag−ATP (APO) complex. **a,** Workflow schematics for cryo-EM data collection, processing, and modelling. Flows continue from left hand side of following row unless otherwise indicated **b,** Gold-standard Fourier shell correlation and Gunier plots **c,** Angular distribution of projections. Resolution is estimated using the 0.143 criterion. **d,** Map anisotropy analysis computed by 3DFSC^1^. **e,** Two views of the cryo-EM map colored by local resolution.

**Extended Data Figure 3.**
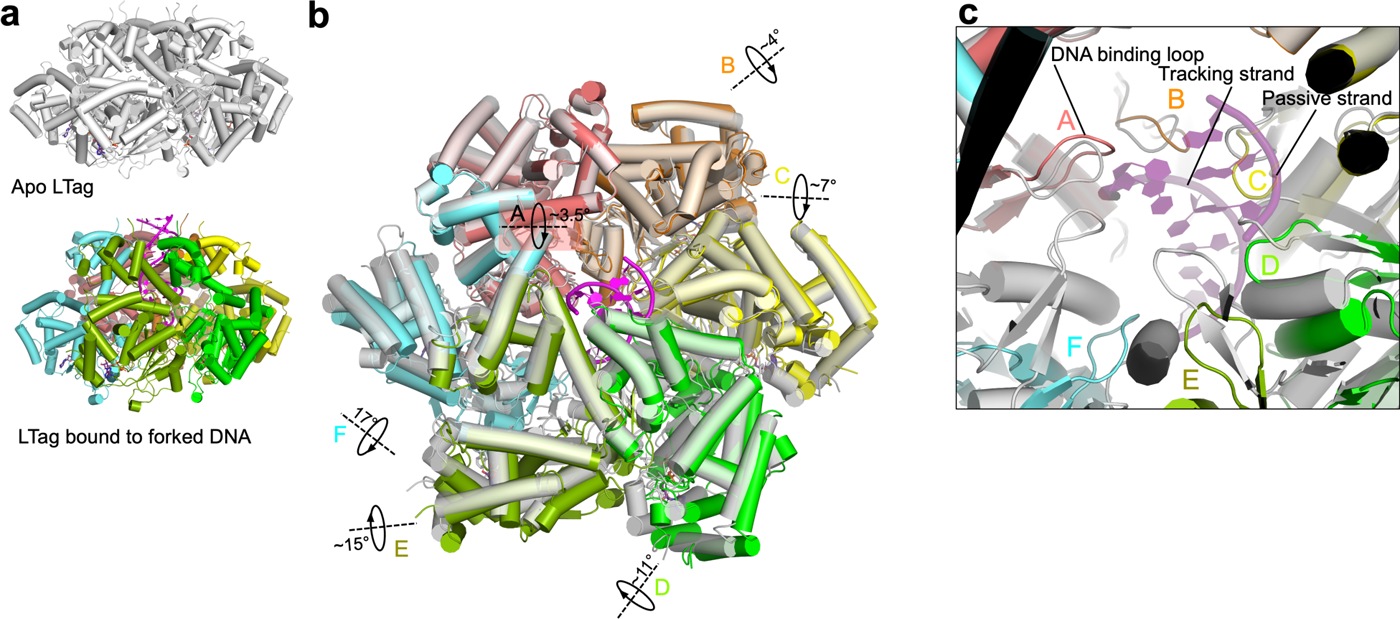
Structural comparison of apo LTag and LTag bound to forked DNA. **a,** Comparative cryo-EM models of the apo (unbound) LTag and LTag bound to forked DNA, **b,** Superposition of the models from **(a)**, illustrating the incremental rotations of the LTag C-tier domains upon DNA binding, attributed to the engagement of DNA-binding loops with the tracking strand. **c,** Close-up view of the LTag central pore in the superimposed structures from **(b)**, comparing the DNA-binding loops. In the apo LTag, loops are shown in an equiplanar arrangement, contrasting with a staircase configuration in the DNA-bound LTag, indicating significant structural reorganization upon DNA engagement.

**Extended Data Figure 4.**
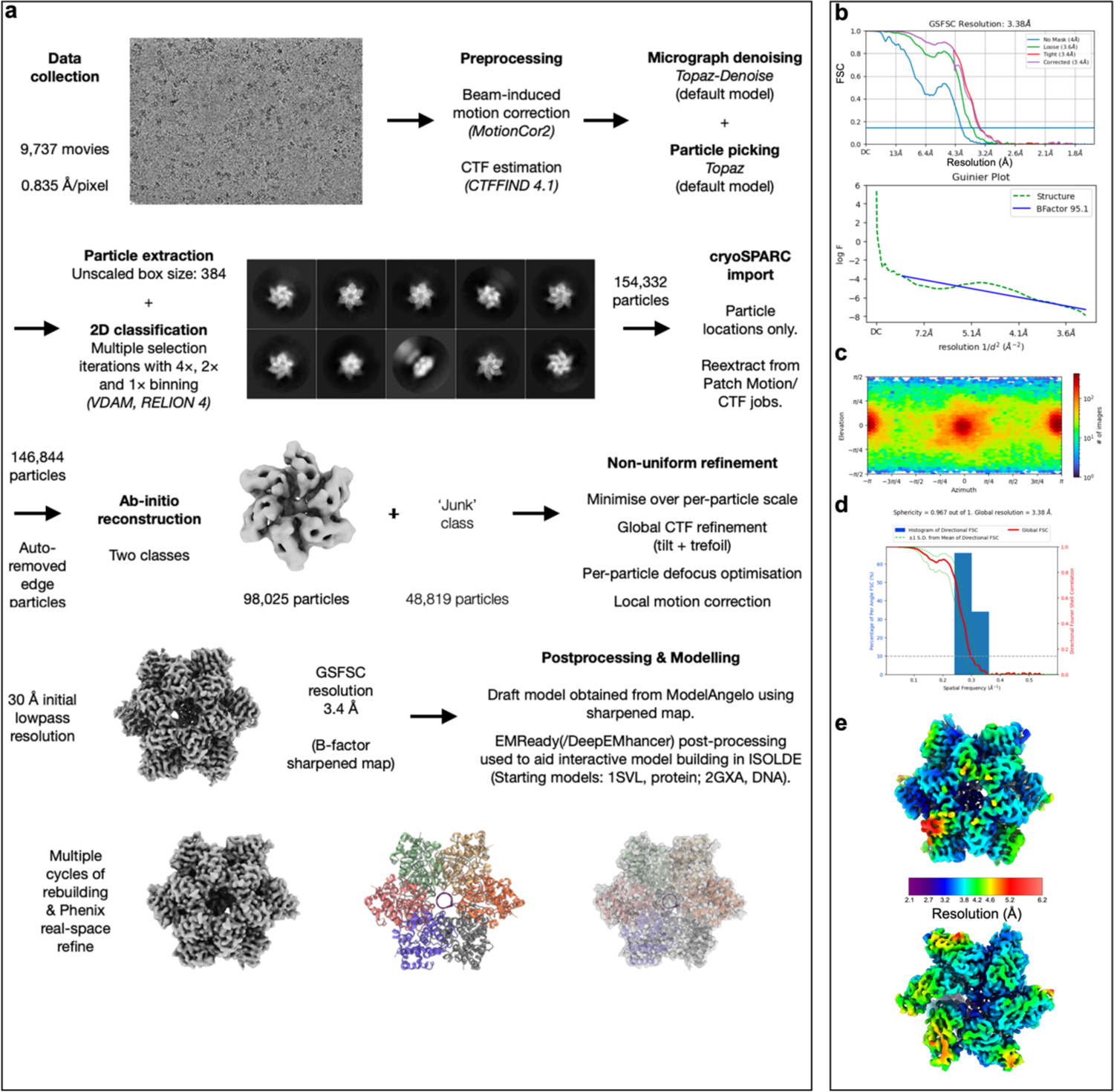
Cryo-EM of the LTag−ADP−DNA complex. **a,** Workflow schematics for cryo-EM data collection, processing, and modelling. Flows continue from left hand side of following row unless otherwise indicated. **b,** Gold-standard Fourier shell correlation and Gunier plots **c,** Angular distribution of projections. Resolution is estimated using the 0.143 criterion. **d,** Map anisotropy analysis computed by 3DFSC^1^. **e,** Two views of the cryo-EM map colored by local resolution.

**Extended Data Figure 5.**
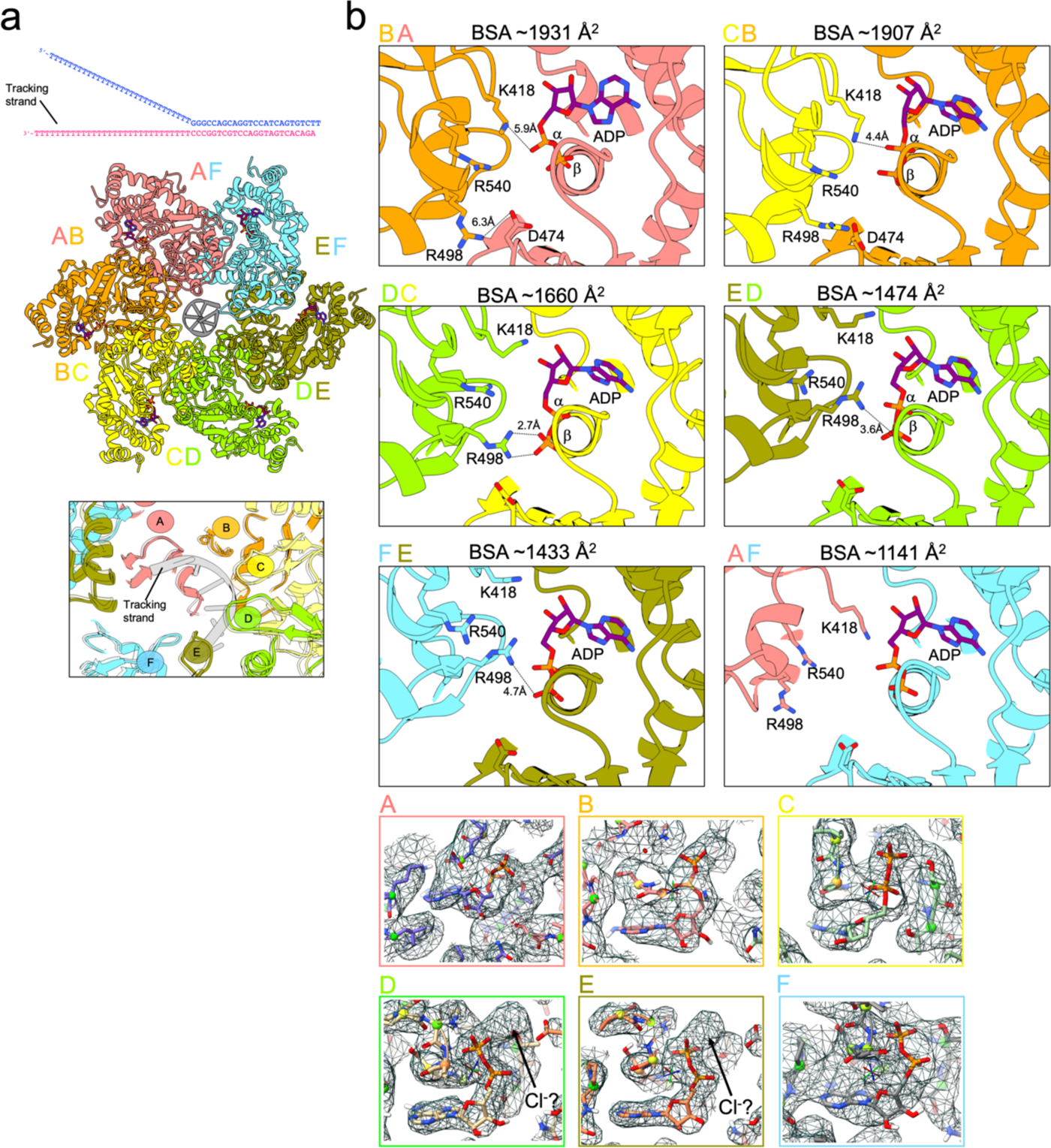
Inter-subunit interactions and nucleotide binding in the LTag−ADP−DNA complex. **a,** Bottom view of the cryo-EM model. The sequence of the DNA substrate is shown above. The lower inset shows details of the DNA binding loop staircase and DNA, with the aligned structure of the LTag*−*ATP*−*DNA complex shown in transparency for comparison. **b,** Inter-subunit interactions at the nucleotide binding pocket. Calculated Buried Surface Areas (BSA) for each interface are shown, along with the assigned interface type. Hydrogen bonds are shown as black dotted lines. **c,** Map density around the ATP molecules in the various LTag subunits. Density at ψ-phosphate position in subunits D and E may be attributed to the presence of a chloride atom.

**Extended Data Figure 6.**
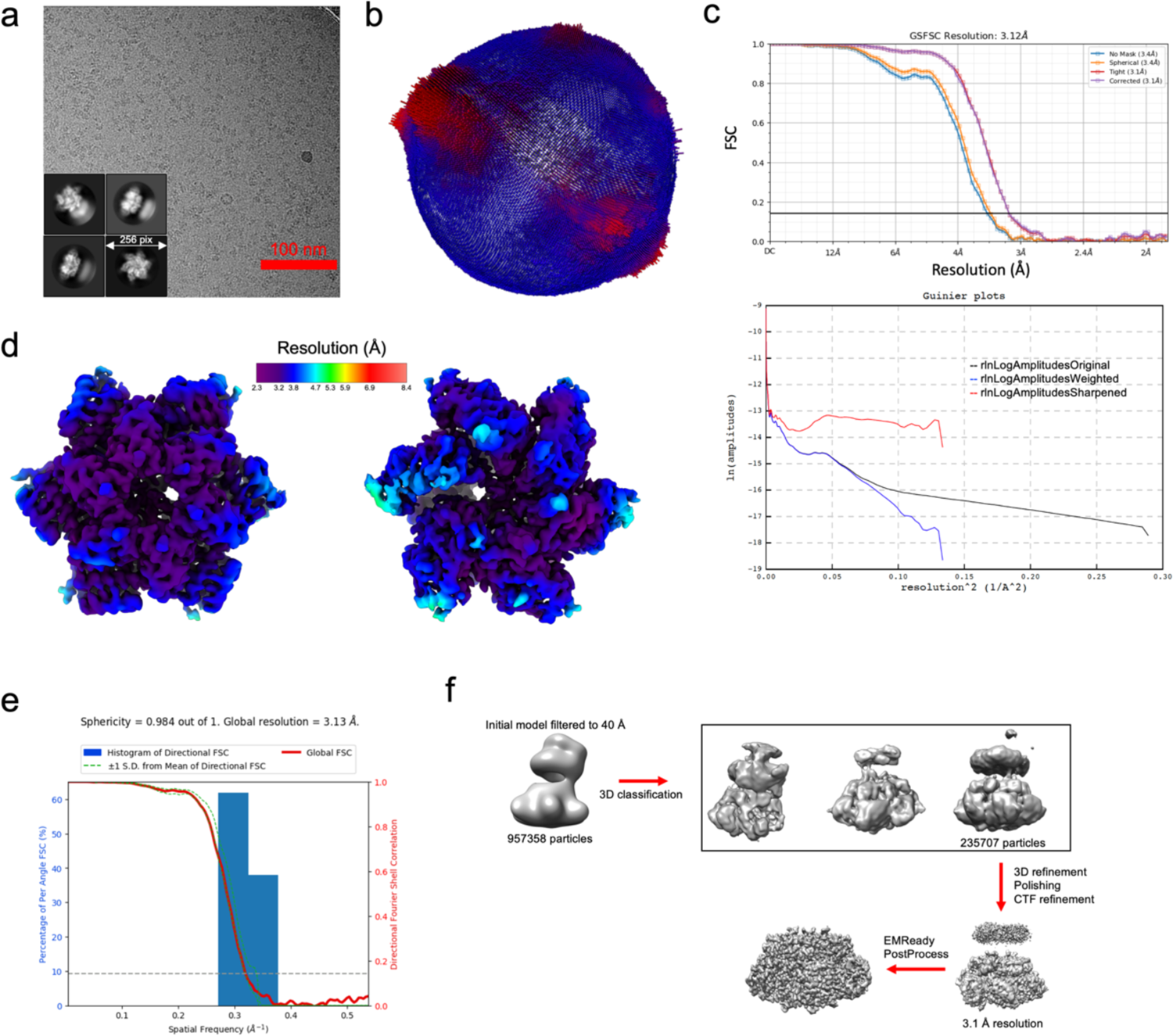
Cryo-EM of the LTag−ANPPNP/Mg^2+^−DNA complex. **a,** Electron micrograph (aligned sum) acquired using a Falcon 4 direct electron detector and representative 2D class averages. **b,** Angular distribution of projections. **c,** Gold-standard Fourier shell correlation and Gunier plots. Resolution is estimated using the 0.143 criterion. **d,** Two views of the cryo-EM map colored by local resolution. **e,** Map anisotropy analysis computed by 3DFSC^1^. **d,** Overview of image processing.

**Extended Data Figure 7.**
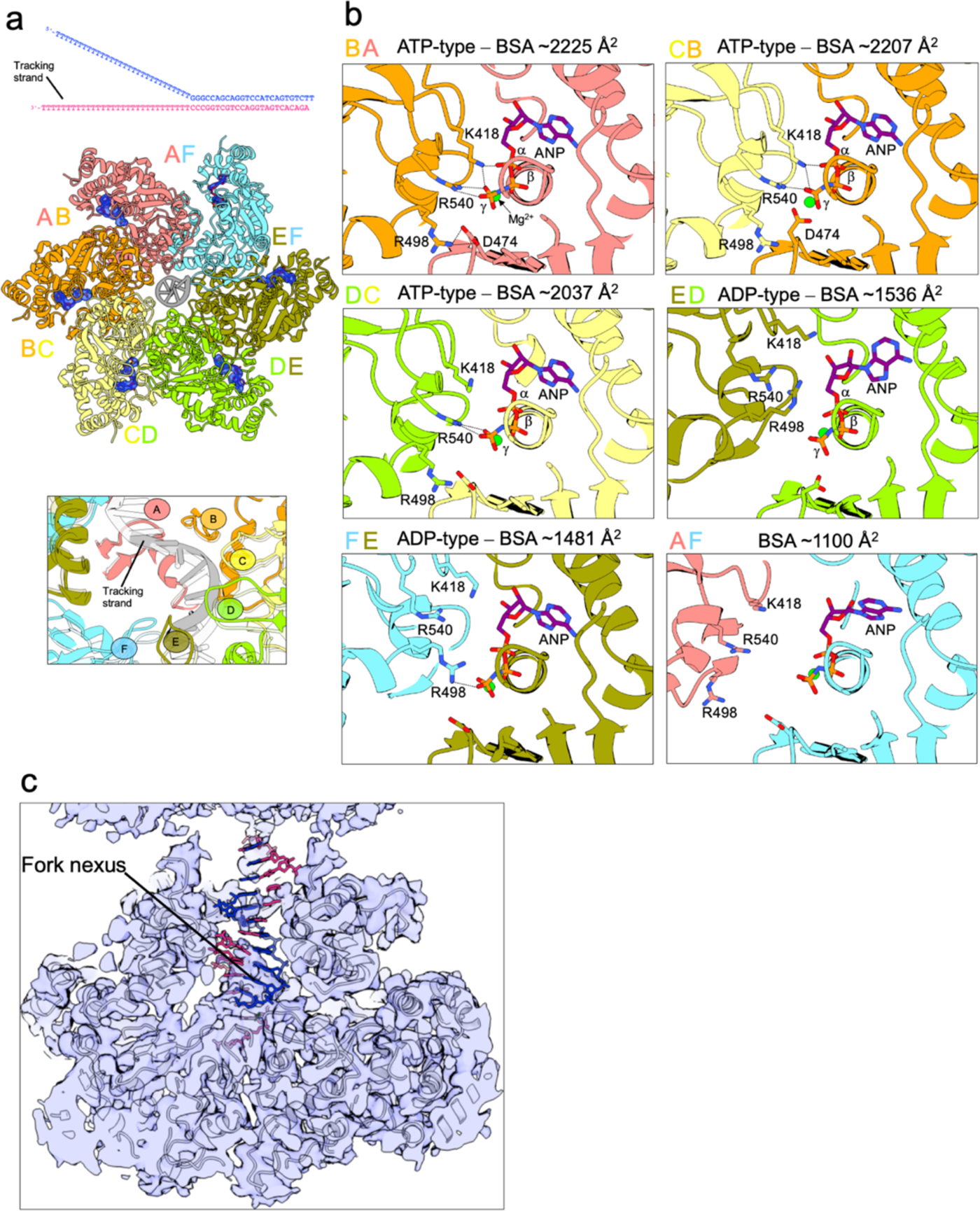
Inter-subunit interactions and nucleotide binding in the LTag−AMP/PNP−DNA complex. **a,** Bottom view of the cryo-EM model. The sequence of the DNA substrate is shown above. The lower inset shows details of the DNA binding loop staircase and DNA. The DNA shown in transparency corresponds to the position in the LTag structure reconstituted with ATP. **b,** Analysis of the nucleotide-binding pocket shows inter-subunit interactions with calculated Buried Surface Areas (BSA) and hydrogen bonds (black dotted lines). **c,** A cut-through of the cryo-EM map displays density at the DNA nexus, with the DNA model (in stick form) aligning with the structure of LTag bound to forked DNA and ATP.

**Extended Data Figure 8.**
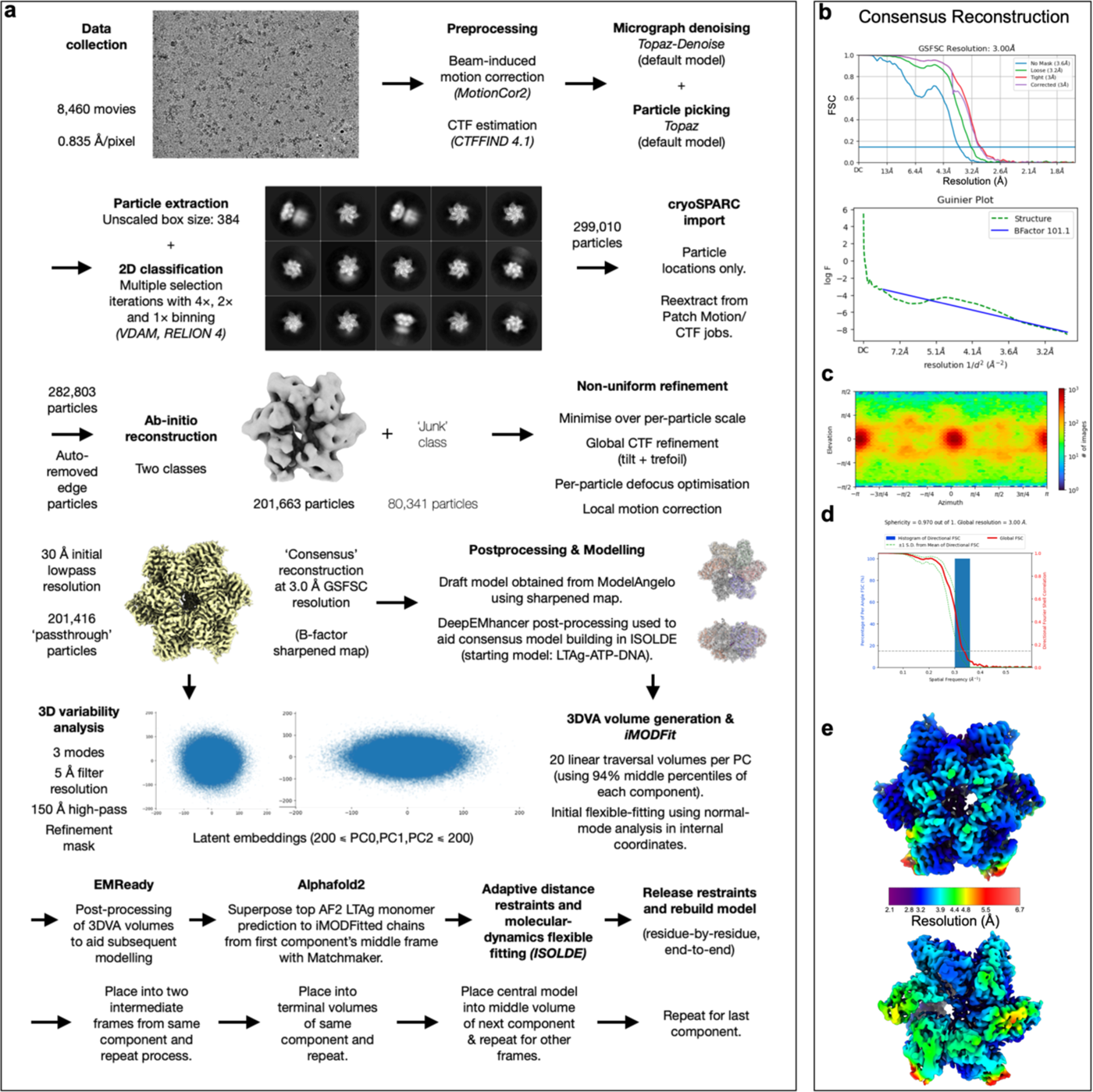
Cryo-EM of the LTag−ATP/Mg^2+^−DNA complex. **a,** Workflow schematics for cryo-EM data collection, processing, and modelling. Flows continue from left hand side of following row unless otherwise indicated. **b,** Gold-standard Fourier shell correlation and Gunier plots for the consensus reconstruction **c,** Angular distribution of projections. Resolution is estimated using the 0.143 criterion. **d,** Map anisotropy analysis computed by 3DFSC^1^. **e,** Two views of the cryo-EM map colored by local resolution.

**Extended Data Figure 9.**
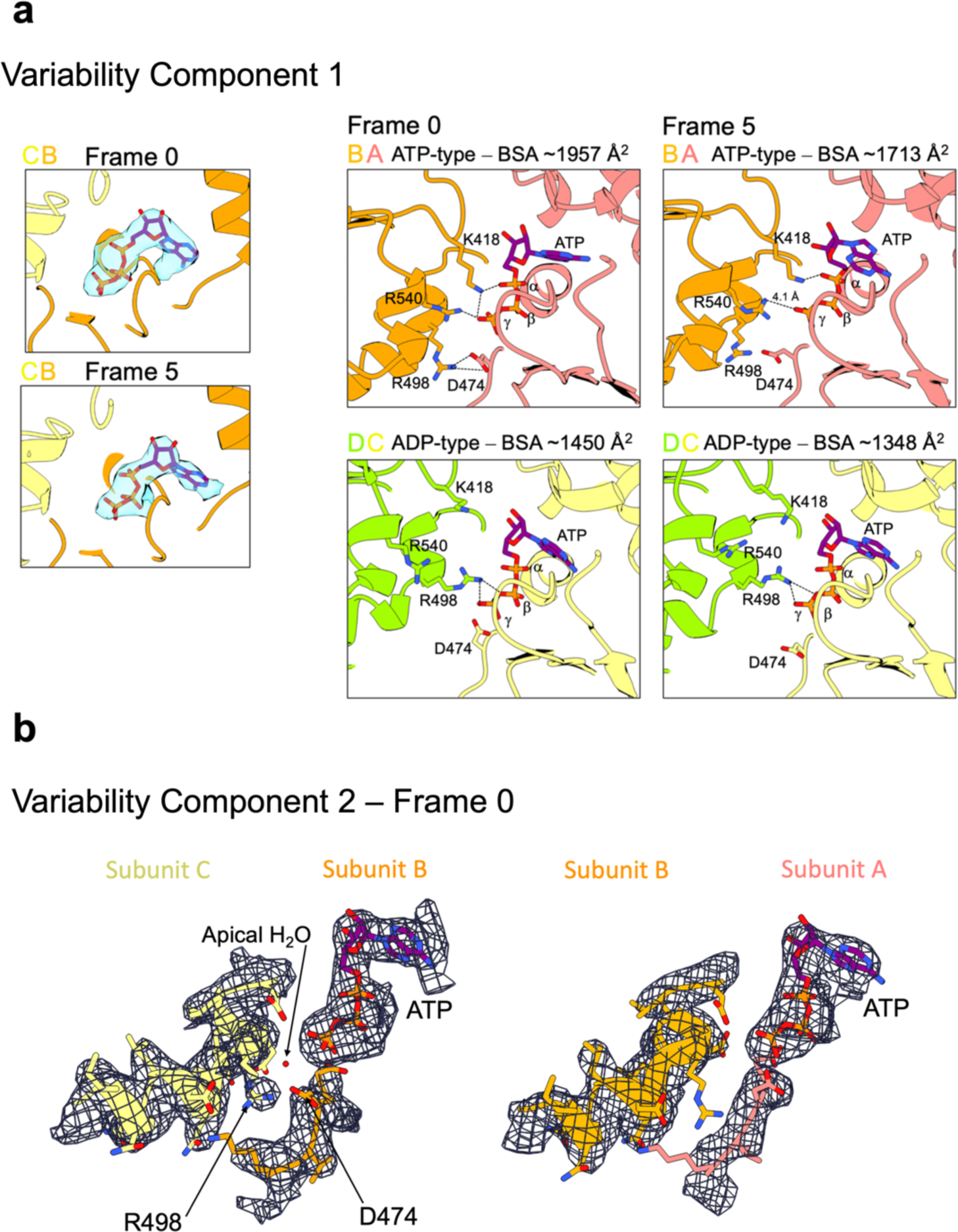
Analysis of the nucleotide binding pocket in the first and second variability trajectories of LTag translocating on forked DNA. **a,** Left: Map density around the ATP molecule at the B/C interface in the initial and final frames of the first component. Density at ψ-phosphate position persists throughout the trajectory despite a full breakage of the interface, suggesting that the hydrolysed phosphate may be retained in the pocket post-hydrolysis. *Right:* Interactions at the nucleotide binding pocket between the A and B or C and D subunits in the initial and final frames of the first variability component. Calculated Buried Surface Areas (BSA) are shown, along with the assigned interface type. Hydrogen bonds are shown as black dotted lines. The analysis reveals a moderate disruption of the A/B interface and no disruption of the C/D interface. **b,** Maps and models of BC and AB interfaces at the initial frame of the second component, showing order of the R498 side chain in C but not B subunits. The position of the apical water molecule, inferred from LTag-ATP crystal structure (PDB ID 1SVM)^2^, is indicated.

**Extended Data Figure 10.**
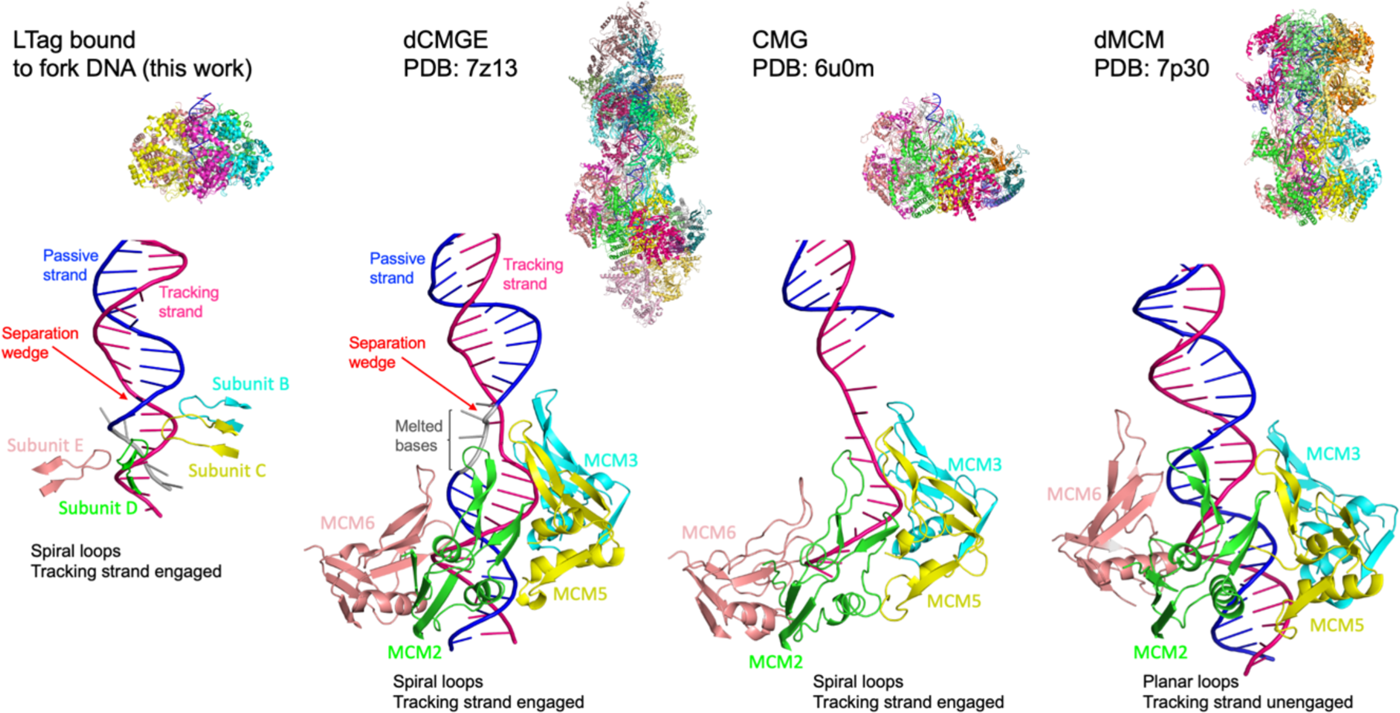
Comparison of internal melting mechanisms in LTag and CMG Helicases. This figure highlights the internal DNA separation mechanism within the LTag helicase bound to forked DNA and its resemblance to the yeast CMG double hexamer structure during DNA melting at the origin^3^ (dCMGE; PDB 7z13). In both cases, the formation of a separation wedge is facilitated by ATP binding, the staircase arrangement of DNA binding loops, and the engagement of these loops with the tracking strand. These elements collectively drive strand separation within the helicase inner chamber, positioning the helicase for ATP-powered translocation. This configuration is mirrored in the actively translocating CMG structure^4^ (PDB 6u0m), showcasing the conserved mechanism of action. Conversely, the pre-CMG formation MCM double hexamer^5^ (dMCM; PDB 7p30) displays planar DNA binding loops and a disengaged tracking strand, indicative of a state not yet primed for ATP-dependent movement.

**Extended Data Table 1.**
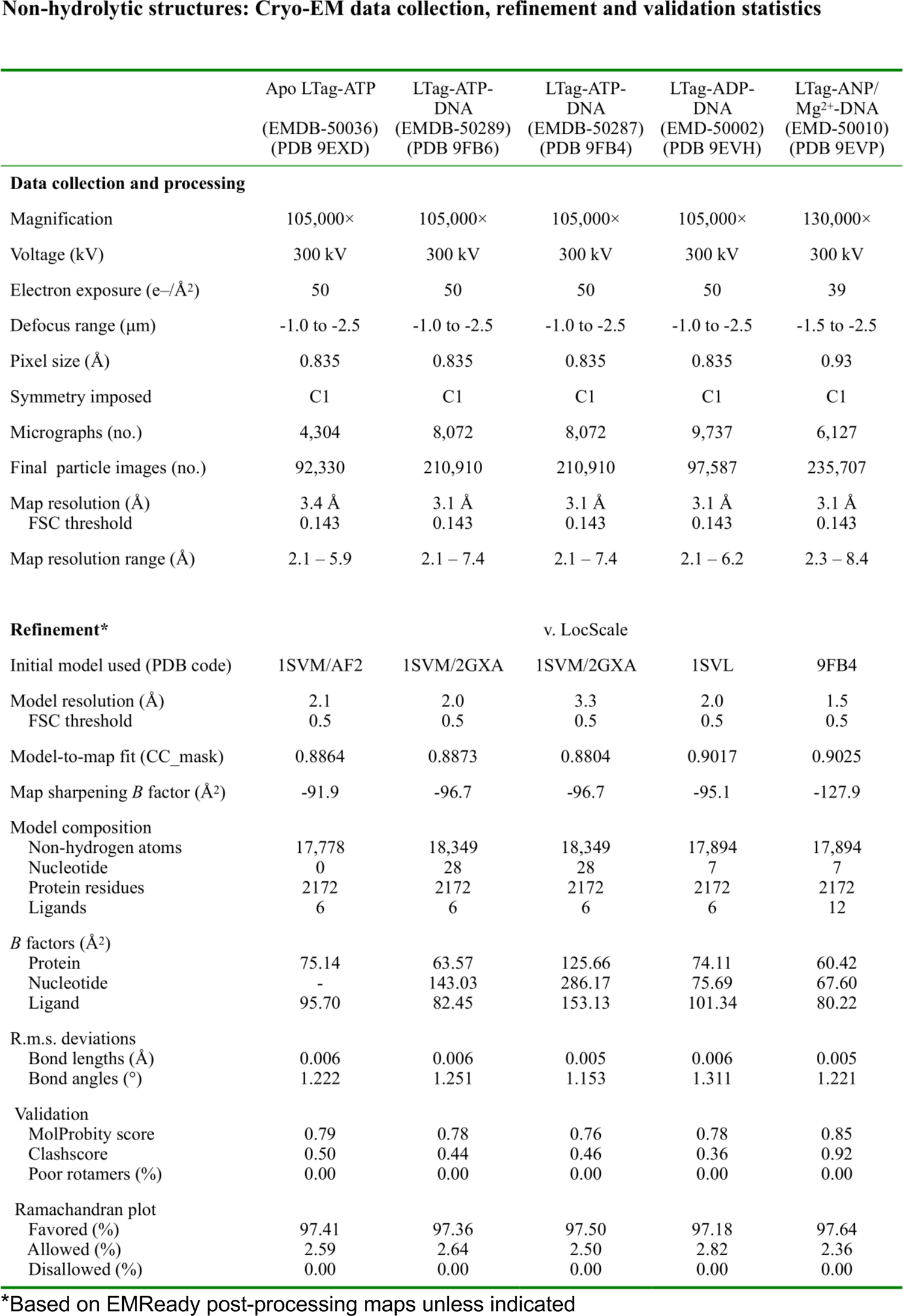

**Extended Data Table 2.**

Component 1 (component\_000): Cryo-EM data collection, refinement and validation statistics
|  | Frame 1<br>(frame_019)<br>(EMDB-50246)<br>(PDB 9F75) | Frame 2<br>(frame_015)<br>(EMDB-50245)<br>(PDB 9F74) | Frame 3<br>(frame_010)<br>(EMD-50179)<br>(PDB 9F3T) | Frame 4<br>(frame_005)<br>(EMD-50190)<br>(PDB 9F5I) | Frame 5<br>(frame_000)<br>(EMD-50250)<br>(PDB 9F7N) |
| --- | --- | --- | --- | --- | --- |
| Data collection and processing |  |  | LTag-ATP/Mg<sup>2+</sup>-DNA |  |  |
| Magnification | 105,000× |  |  |  |  |
| Voltage (kV) | 300 kV |  |  |  |  |
| Electron exposure (e-/Å<sup>2</sup>) | 50 e-/Å<sup>2</sup> |  |  |  |  |
| Defocus range (μm) | -1.0 to -2.5 μm |  |  |  |  |
| Pixel size (Å) | 0.835 Å/pixel |  |  |  |  |
| Symmetry imposed | C1 |  |  |  |  |
| Micrographs (no.) | 8,460 |  |  |  |  |
| Final particle images (no.) | 201,416 |  |  |  |  |
| Map resolution (Å) | 3.00 Å (Consensus reconstruction, EMD-50290) |  |  |  |  |
| FSC threshold | 0.143 |  |  |  |  |
| Map resolution range (Å) | 2.1 – 6.7 |  |  |  |  |
| Refinement* |  |  |  |  |  |
| Initial model used (PDB code) | 9FB4, AlphaFold2 |  |  |  |  |
| Model resolution (Å)* | 2.2 | 2.1 | 2.1 | 2.2 | 2.2 |
| FSC threshold | 0.5 | 0.5 | 0.5 | 0.5 | 0.5 |
| Model-to-map fit (CC_mask) | 0.8971 | 0.9010 | 0.8997 | 0.8953 | 0.8878 |
| Map sharpening B factor (Å<sup>2</sup>) | -101.1 (Consensus reconstruction, EMD-50290) |  |  |  |  |
| Model composition |  |  |  |  |  |
| Non-hydrogen atoms | 17,938 |  |  |  |  |
| Nucleotide | 8 |  |  |  |  |
| Protein residues | 2172 |  |  |  |  |
| Ligands | 6 |  |  |  |  |
| B factors (Å<sup>2</sup>) |  |  |  |  |  |
| Protein | 101.80 | 100.24 | 107.27 | 108.30 | 109.02 |
| Nucleotide | 130.45 | 116.14 | 107.27 | 165.57 | 168.31 |
| Ligand | 123.45 | 122.17 | 130.12 | 135.53 | 139.05 |
| R.m.s. deviations |  |  |  |  |  |
| Bond lengths (Å) | 0.007 | 0.007 | 0.007 | 0.007 | 0.007 |
| Bond angles (°) | 1.193 | 1.155 | 1.163 | 1.159 | 1.180 |
| Validation |  |  |  |  |  |
| MolProbity score | 1.22 | 1.44 | 1.06 | 1.00 | 1.23 |
| Clashscore | 1.86 | 1.06 | 1.28 | 1.42 | 1.94 |
| Poor rotamers (%) | 0.00 | 0.00 | 0.00 | 0.00 | 0.00 |
| Ramachandran plot |  |  |  |  |  |
| Favored (%) | 96.06 | 96.94 | 96.67 | 97.36 | 96.06 |
| Allowed (%) | 3.94 | 3.06 | 3.33 | 2.64 | 3.94 |
| Disallowed (%) | 0.00 | 0.00 | 0.00 | 0.00 | 0.00 |

\*Based on EMReady post-processing maps unless indicated

**Extended Data Table 3.**

Component 2 (component\_001): Cryo-EM data collection, refinement and validation statistics
|  | Frame 1<br>(frame_019)<br>(EMDB-50261)<br>(PDB 9F9W) | Frame 2<br>(frame_015)<br>(EMDB-50257)<br>(PDB 9F9O) | Frame 3<br>(frame_010)<br>(EMD-50180)<br>(PDB 9F3U) | Frame 4<br>(frame_005)<br>(EMD-50256)<br>(PDB 9F9N) | Frame 5<br>(frame_000)<br>(EMD-50262)<br>(PDB 9F9X) |
| --- | --- | --- | --- | --- | --- |
| **Data collection and processing** | LTag-ATP/Mg<sup>2+</sup>-DNA |  |  |  |  |
| Magnification | 105,000× |  |  |  |  |
| Voltage (kV) | 300 kV |  |  |  |  |
| Electron exposure (e-/Å<sup>2</sup>) | 50 e-/Å<sup>2</sup> |  |  |  |  |
| Defocus range (μm) | -1.0 to -2.5 μm |  |  |  |  |
| Pixel size (Å) | 0.835 Å/pixel |  |  |  |  |
| Symmetry imposed | C1 |  |  |  |  |
| Micrographs (no.) | 8,460 |  |  |  |  |
| Final particle images (no.) | 201,416 |  |  |  |  |
| Map resolution (Å) | 3.00 Å (Consensus reconstruction, EMD-50290) |  |  |  |  |
| FSC threshold | 0.143 |  |  |  |  |
| Map resolution range (Å) | 2.1 – 6.7 |  |  |  |  |
| **Refinement** |  |  |  |  |  |
| Initial model used (PDB code) | 9FB4, 9F3T, AlphaFold2 |  |  |  |  |
| Model resolution (Å)* | 2.3 | 2.2 | 2.2 | 2.2 | 2.2 |
| FSC threshold | 0.5 | 0.5 | 0.5 | 0.5 | 0.5 |
| Model-to-map fit (CC_mask) | 0.8865 | 0.8972 | 0.8995 | 0.8941 | 0.8851 |
| Map sharpening *B* factor (Å<sup>2</sup>) | -101.1 (Consensus reconstruction, EMD-50290) |  |  |  |  |
| Model composition |  |  |  |  |  |
| Non-hydrogen atoms | 17,938 |  |  |  |  |
| Nucleotide | 8 |  |  |  |  |
| Protein residues | 2172 |  |  |  |  |
| Ligands | 6 |  |  |  |  |
| *B* factors (Å<sup>2</sup>) |  |  |  |  |  |
| Protein | 104.12 | 103.25 | 106.18 | 109.45 | 110.49 |
| Nucleotide | 112.14 | 124.56 | 161.07 | 158.15 | 167.99 |
| Ligand | 127.72 | 132.28 | 128.79 | 128.46 | 126.27 |
| R.m.s. deviations |  |  |  |  |  |
| Bond lengths (Å) | 0.007 | 0.007 | 0.007 | 0.007 | 0.007 |
| Bond angles (°) | 1.201 | 1.164 | 1.158 | 1.166 | 1.188 |
| Validation |  |  |  |  |  |
| MolProbity score | 1.10 | 1.06 | 1.05 | 1.05 | 1.18 |
| Clashscore | 1.44 | 1.33 | 1.36 | 1.44 | 1.86 |
| Poor rotamers (%) | 0.00 | 0.00 | 0.00 | 0.00 | 0.00 |
| Ramachandran plot |  |  |  |  |  |
| Favored (%) | 96.57 | 96.76 | 96.94 | 97.04 | 96.44 |
| Allowed (%) | 3.43 | 3.24 | 3.06 | 2.96 | 3.56 |
| Disallowed (%) | 0.00 | 0.00 | 0.00 | 0.00 | 0.00 |

\*Based on EMReady post-processing maps unless indicated

**Extended Data Table 4.**

|  | Frame 1<br>(frame_000)<br>(EMDB-50288)<br>(PDB 9FB5) | Frame 2<br>(frame_005)<br>(EMDB-50265)<br>(PDB 9FA2) | Frame 3<br>(frame_010)<br>(EMD-50264)<br>(PDB 9FA1) | Frame 4<br>(frame_015)<br>(EMD-50244)<br>(PDB 9F73) | Frame 5<br>(frame_019)<br>(EMD-50286)<br>(PDB 9FB0) |
| --- | --- | --- | --- | --- | --- |
| **Data collection and processing** | LTag-ATP/Mg<sup>2+</sup>-DNA |  |  |  |  |
| Magnification | 105,000× |  |  |  |  |
| Voltage (kV) | 300 kV |  |  |  |  |
| Electron exposure (e-/Å<sup>2</sup>) | 50 e-/Å<sup>2</sup> |  |  |  |  |
| Defocus range (μm) | -1.0 to -2.5 μm |  |  |  |  |
| Pixel size (Å) | 0.835 Å/pixel |  |  |  |  |
| Symmetry imposed | C1 |  |  |  |  |
| Micrographs (no.) | 8,460 |  |  |  |  |
| Final particle images (no.) | 201,416 |  |  |  |  |
| Map resolution (Å) | 3.00 Å (Consensus reconstruction, EMD-50290) |  |  |  |  |
| FSC threshold | 0.143 |  |  |  |  |
| Map resolution range (Å) | 2.1 – 6.7 |  |  |  |  |
| **Refinement\*** |  |  |  |  |  |
| Initial model used (PDB code) | 9FB4, 9F3T, 9F3U, AlphaFold2 |  |  |  |  |
| Model resolution (Å)* | 2.3 | 2.2 | 2.1 | 2.2 | 2.2 |
| FSC threshold | 0.5 | 0.5 | 0.5 | 0.5 | 0.5 |
| Model-to-map fit (CC_mask) | 0.8766 | 0.8927 | 0.9004 | 0.8939 | 0.8864 |
| Map sharpening *B* factor (Å<sup>2</sup>) | -101.1 (Consensus reconstruction, EMD-50290) |  |  |  |  |
| Model composition |  |  |  |  |  |
| Non-hydrogen atoms | 17,938 |  |  |  |  |
| Nucleotide | 8 |  |  |  |  |
| Protein residues | 2172 |  |  |  |  |
| Ligands | 6 |  |  |  |  |
| *B* factors (Å<sup>2</sup>) |  |  |  |  |  |
| Protein | 113.59 | 110.76 | 108.34 | 111.22 | 113.56 |
| Nucleotide | 126.71 | 163.35 | 182.28 | 127.58 | 160.10 |
| Ligand | 129.54 | 133.02 | 131.09 | 132.11 | 131.56 |
| R.m.s. deviations |  |  |  |  |  |
| Bond lengths (Å) | 0.007 | 0.007 | 0.007 | 0.007 | 0.007 |
| Bond angles (°) | 1.217 | 1.182 | 1.151 | 1.193 | 1.208 |
| Validation |  |  |  |  |  |
| MolProbity score | 1.44 | 1.14 | 1.02 | 1.15 | 1.23 |
| Clashscore | 2.89 | 1.42 | 1.22 | 1.47 | 1.72 |
| Poor rotamers (%) | 0.00 | 0.00 | 0.00 | 0.00 | 0.00 |
| Ramachandran plot |  |  |  |  |  |
| Favored (%) | 94.77 | 96.11 | 96.94 | 96.06 | 95.69 |
| Allowed (%) | 5.23 | 3.89 | 3.06 | 3.94 | 4.31 |
| Disallowed (%) | 0.00 | 0.00 | 0.00 | 0.00 | 0.00 |

\*Based on EMReady post-processing maps unless indicated

**Extended Data Table 5.**
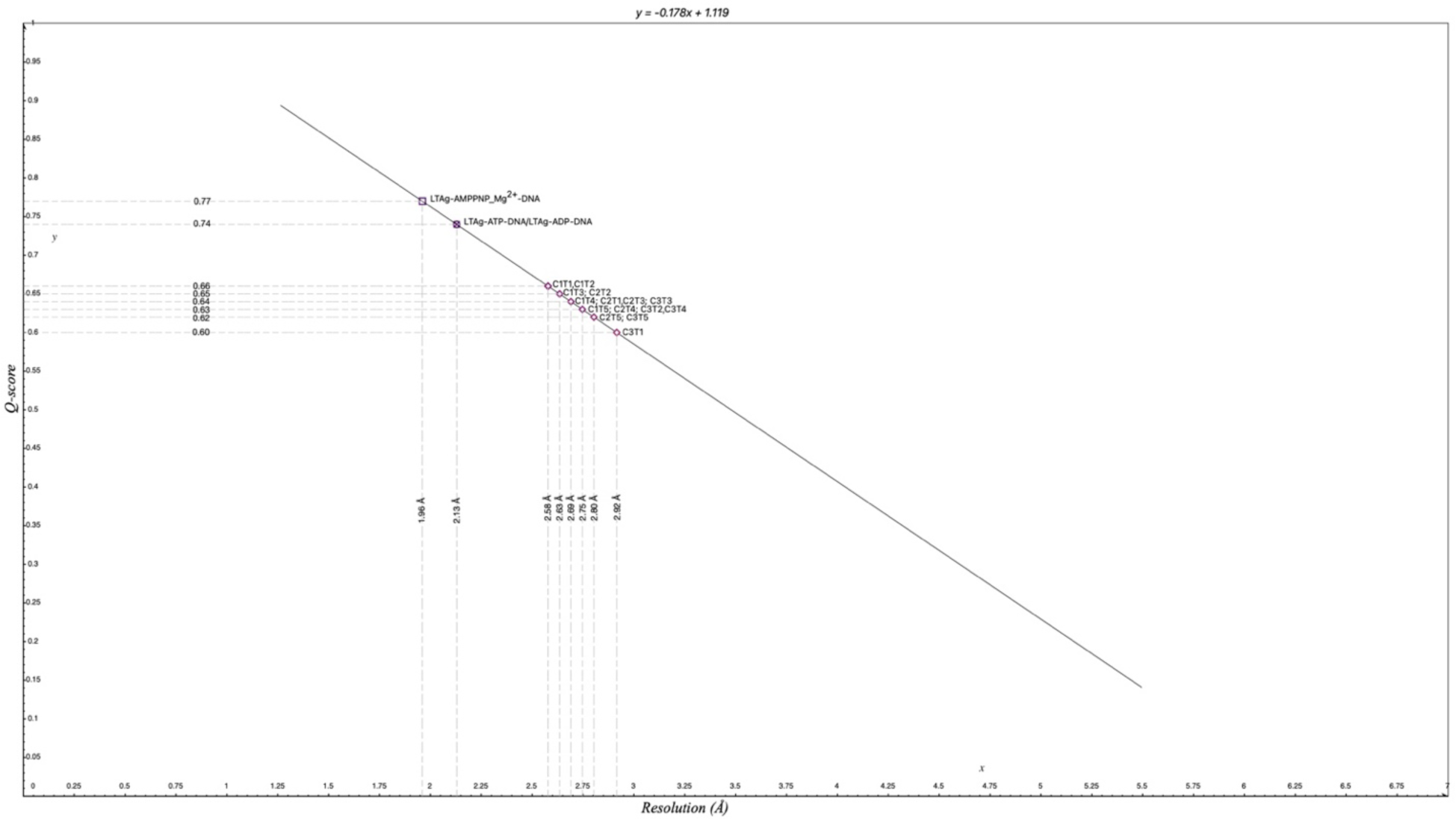
Q-scores^6^ of modelled structures against respective maps postprocessed by EMReady^7^. (horizontal dotted lines). C1, C2, C3 refer to 3DVA principal components. T1, T2, T3, T4, T5 refer to modelled volumes from each component, in order of reaction sequence. Vertical dotted lines indicate GSFSC resolutions ordinarily corresponding to said Q-scores, underscoring quality of modelling. Solid line represents linear fit of average Q-score versus reported resolution of deposited map-model pairs in EMDB/PDB^6^.

## References

1 Fernandez, A. J. & Berger, J. M. Mechanisms of hexameric helicases. Crit Rev Biochem Mol Biol 56, 621–639 (2021). 10.1080/10409238.2021.1954597

2 Gao, Y. & Yang, W. Different mechanisms for translocation by monomeric and hexameric helicases. Curr Opin Struct Biol 61, 25–32 (2020). 10.1016/j.sbi.2019.10.003

3 Fanning, E. Simian virus 40 large T antigen: the puzzle, the pieces, and the emerging picture. J Virol 66, 1289–1293 (1992). 10.1128/JVI.66.3.1289-1293.1992

4 Stillman, B. Smart machines at the DNA replication fork. Cell 78, 725–728 (1994). 10.1016/s0092-8674(94)90362-x

5 Waga, S. & Stillman, B. The DNA replication fork in eukaryotic cells. Annu Rev Biochem 67, 721–751 (1998). 10.1146/annurev.biochem.67.1.721

6 Douglas, M. E., Ali, F. A., Costa, A. & Diffley, J. F. X. The mechanism of eukaryotic CMG helicase activation. Nature 555, 265–268 (2018). 10.1038/nature25787

7 Ilves, I., Petojevic, T., Pesavento, J. J. & Botchan, M. R. Activation of the MCM2-7 helicase by association with Cdc45 and GINS proteins. Mol Cell 37, 247–258 (2010). 10.1016/j.molcel.2009.12.030

8 Moyer, S. E., Lewis, P. W. & Botchan, M. R. Isolation of the Cdc45/Mcm2-7/GINS (CMG) complex, a candidate for the eukaryotic DNA replication fork helicase. Proc Natl Acad Sci U S A 103, 10236–10241 (2006). 10.1073/pnas.0602400103

9 Chang, Y. P. et al. Mechanism of origin DNA recognition and assembly of an initiator-helicase complex by SV40 large tumor antigen. Cell Rep 3, 1117–1127 (2013). 10.1016/j.celrep.2013.03.002

10 Gai, D., Zhao, R., Li, D., Finkielstein, C. V. & Chen, X. S. Mechanisms of conformational change for a replicative hexameric helicase of SV40 large tumor antigen. Cell 119, 47–60 (2004). 10.1016/j.cell.2004.09.017

11 Li, D. et al. Structure of the replicative helicase of the oncoprotein SV40 large tumour antigen. Nature 423, 512–518 (2003). 10.1038/nature01691

12 Luo, X., Sanford, D. G., Bullock, P. A. & Bachovchin, W. W. Solution structure of the origin DNA-binding domain of SV40 T-antigen. Nat Struct Biol 3, 1034–1039 (1996). 10.1038/nsb1296-1034

13 Cuesta, I. et al. Conformational rearrangements of SV40 large T antigen during early replication events. J Mol Biol 397, 1276–1286 (2010). 10.1016/j.jmb.2010.02.042

14 Dean, F. B., Borowiec, J. A., Eki, T. & Hurwitz, J. The simian virus 40 T antigen double hexamer assembles around the DNA at the replication origin. J Biol Chem 267, 14129–14137 (1992).

15 Mastrangelo, I. A. et al. ATP-dependent assembly of double hexamers of SV40 T antigen at the viral origin of DNA replication. Nature 338, 658–662 (1989). 10.1038/338658a0

16 Parsons, R. E. et al. Cooperative assembly of simian virus 40 T-antigen hexamers on functional halves of the replication origin. J Virol 65, 2798–2806 (1991). 10.1128/JVI.65.6.2798-2806.1991

17 Tjian, R. The binding site on SV40 DNA for a T antigen-related protein. Cell 13, 165–179 (1978). 10.1016/0092-8674(78)90147-2

18 Langston, L. D., Yuan, Z., Georgescu, R., Li, H. & O’Donnell, M. E. SV40 T-antigen uses a DNA shearing mechanism to initiate origin unwinding. Proc Natl Acad Sci U S A 119, e2216240119 (2022). 10.1073/pnas.2216240119

19 Dodson, M., Dean, F. B., Bullock, P., Echols, H. & Hurwitz, J. Unwinding of duplex DNA from the SV40 origin of replication by T antigen. Science 238, 964–967 (1987). 10.1126/science.2823389

20 Yardimci, H. et al. Bypass of a protein barrier by a replicative DNA helicase. Nature 492, 205–209 (2012). 10.1038/nature11730

21 Itsathitphaisarn, O., Wing, R. A., Eliason, W. K., Wang, J. & Steitz, T. A. The hexameric helicase DnaB adopts a nonplanar conformation during translocation. Cell 151, 267–277 (2012). 10.1016/j.cell.2012.09.014

22 Rzechorzek, N. J., Hardwick, S. W., Jatikusumo, V. A., Chirgadze, D. Y. & Pellegrini, L. CryoEM structures of human CMG-ATPgammaS-DNA and CMG-AND-1 complexes. Nucleic Acids Res 48, 6980–6995 (2020). 10.1093/nar/gkaa429

23 Thomsen, N. D. & Berger, J. M. Running in reverse: the structural basis for translocation polarity in hexameric helicases. Cell 139, 523–534 (2009). 10.1016/j.cell.2009.08.043

24 Meagher, M., Epling, L. B. & Enemark, E. J. DNA translocation mechanism of the MCM complex and implications for replication initiation. Nat Commun 10, 3117 (2019). 10.1038/s41467-019-11074-3

25 Enemark, E. J. & Joshua-Tor, L. Mechanism of DNA translocation in a replicative hexameric helicase. Nature 442, 270–275 (2006). 10.1038/nature04943

26 Eickhoff, P. et al. Molecular Basis for ATP-Hydrolysis-Driven DNA Translocation by the CMG Helicase of the Eukaryotic Replisome. Cell Rep 28, 2673–2688 e2678 (2019). 10.1016/j.celrep.2019.07.104

27 Gao, Y. et al. Structures and operating principles of the replisome. Science 363 (2019). 10.1126/science.aav7003

28 Punjani, A. & Fleet, D. J. 3D variability analysis: Resolving continuous flexibility and discrete heterogeneity from single particle cryo-EM. J Struct Biol 213, 107702 (2021). 10.1016/j.jsb.2021.107702

29 He, J., Li, T. & Huang, S. Y. Improvement of cryo-EM maps by simultaneous local and non-local deep learning. Nat Commun 14, 3217 (2023). 10.1038/s41467-023-39031-1

30 Croll, T. I. ISOLDE: a physically realistic environment for model building into low-resolution electron-density maps. Acta Crystallogr D Struct Biol 74, 519–530 (2018). 10.1107/S2059798318002425

31 Pintilie, G. et al. Measurement of atom resolvability in cryo-EM maps with Q-scores. Nat Methods 17, 328–334 (2020). 10.1038/s41592-020-0731-1

32 Shein, M. et al. Characterizing ATP processing by the AAA+ protein p97 at the atomic level. Nat Chem (2024). 10.1038/s41557-024-01440-0

33 Yuan, Z. et al. Structure of the eukaryotic replicative CMG helicase suggests a pumpjack motion for translocation. Nat Struct Mol Biol 23, 217–224 (2016). 10.1038/nsmb.3170

34 Pellegrini, L. The CMG DNA helicase and the core replisome. Curr Opin Struct Biol 81, 102612 (2023). 10.1016/j.sbi.2023.102612

35 Xu, Z. et al. Synergism between CMG helicase and leading strand DNA polymerase at replication fork. Nat Commun 14, 5849 (2023). 10.1038/s41467-023-41506-0

36 Abramson, J. et al. Accurate structure prediction of biomolecular interactions with AlphaFold 3. Nature (2024). 10.1038/s41586-024-07487-w

37 Fei, X., Bell, T. A., Barkow, S. R., Baker, T. A. & Sauer, R. T. Structural basis of ClpXP recognition and unfolding of ssrA-tagged substrates. Elife 9 (2020). 10.7554/eLife.61496

38 Ripstein, Z. A., Huang, R., Augustyniak, R., Kay, L. E. & Rubinstein, J. L. Structure of a AAA+ unfoldase in the process of unfolding substrate. Elife 6 (2017). 10.7554/eLife.25754

39 Borowiec, J. A. & Hurwitz, J. Localized melting and structural changes in the SV40 origin of replication induced by T-antigen. EMBO J 7, 3149–3158 (1988). 10.1002/j.1460-2075.1988.tb03182.x

40 Wiekowski, M., Schwarz, M. W. & Stahl, H. Simian virus 40 large T antigen DNA helicase. Characterization of the ATPase-dependent DNA unwinding activity and its substrate requirements. J Biol Chem 263, 436–442 (1988).

41 Remus, D. et al. Concerted loading of Mcm2-7 double hexamers around DNA during DNA replication origin licensing. Cell 139, 719–730 (2009). 10.1016/j.cell.2009.10.015

42 Greiwe, J. F. et al. Structural mechanism for the selective phosphorylation of DNA-loaded MCM double hexamers by the Dbf4-dependent kinase. Nat Struct Mol Biol 29, 10–20 (2022). 10.1038/s41594-021-00698-z

43 Lewis, J. S. et al. Mechanism of replication origin melting nucleated by CMG helicase assembly. Nature 606, 1007–1014 (2022). 10.1038/s41586-022-04829-4

44 Yuan, Z. et al. DNA unwinding mechanism of a eukaryotic replicative CMG helicase. Nat Commun 11, 688 (2020). 10.1038/s41467-020-14577-6

45 SenGupta, D. J. & Borowiec, J. A. Strand-specific recognition of a synthetic DNA replication fork by the SV40 large tumor antigen. Science 256, 1656–1661 (1992). 10.1126/science.256.5064.1656

46 Bokori-Brown, M. et al. Cryo-EM structure of lysenin pore elucidates membrane insertion by an aerolysin family protein. Nat Commun 7, 11293 (2016). 10.1038/ncomms11293

47 Kimanius, D., Dong, L., Sharov, G., Nakane, T. & Scheres, S. H. W. New tools for automated cryo-EM single-particle analysis in RELION-4.0. Biochem J 478, 4169–4185 (2021). 10.1042/BCJ20210708

48 Zheng, S. Q. et al. MotionCor2: anisotropic correction of beam-induced motion for improved cryo-electron microscopy. Nat Methods 14, 331–332 (2017). 10.1038/nmeth.4193

49 Rohou, A. & Grigorieff, N. CTFFIND4: Fast and accurate defocus estimation from electron micrographs. J Struct Biol 192, 216–221 (2015). 10.1016/j.jsb.2015.08.008

50 Bepler, T. et al. Positive-unlabeled convolutional neural networks for particle picking in cryo-electron micrographs. Nat Methods 16, 1153–1160 (2019). 10.1038/s41592-019-0575-8

51 Tan, Y. Z. et al. Addressing preferred specimen orientation in single-particle cryo-EM through tilting. Nat Methods 14, 793–796 (2017). 10.1038/nmeth.4347

52 Jakobi, A. J., Wilmanns, M. & Sachse, C. Model-based local density sharpening of cryo-EM maps. Elife 6 (2017). 10.7554/eLife.27131

53 Jumper, J. et al. Highly accurate protein structure prediction with AlphaFold. Nature 596, 583–589 (2021). 10.1038/s41586-021-03819-2

54 Meng, E. C., Pettersen, E. F., Couch, G. S., Huang, C. C. & Ferrin, T. E. Tools for integrated sequence-structure analysis with UCSF Chimera. BMC Bioinformatics 7, 339 (2006). 10.1186/1471-2105-7-339

55 Kabsch, W. & Sander, C. Dictionary of protein secondary structure: pattern recognition of hydrogen-bonded and geometrical features. Biopolymers 22, 2577–2637 (1983). 10.1002/bip.360221211

56 Pettersen, E. F. et al. UCSF Chimera--a visualization system for exploratory research and analysis. J Comput Chem 25, 1605–1612 (2004). 10.1002/jcc.20084

57 Meng, E. C. et al. UCSF ChimeraX: Tools for structure building and analysis. Protein Sci 32, e4792 (2023). 10.1002/pro.4792

## Extended Data References

1 Tan, Y. Z. et al. Addressing preferred specimen orientation in single-particle cryo-EM through tilting. Nat Methods 14, 793–796 (2017). 10.1038/nmeth.4347

2 Gai, D., Zhao, R., Li, D., Finkielstein, C. V. & Chen, X. S. Mechanisms of conformational change for a replicative hexameric helicase of SV40 large tumor antigen. Cell 119, 47–60 (2004). 10.1016/j.cell.2004.09.017

3 Lewis, J. S. et al. Mechanism of replication origin melting nucleated by CMG helicase assembly. Nature 606, 1007–1014 (2022). 10.1038/s41586-022-04829-4

4 Yuan, Z. et al. DNA unwinding mechanism of a eukaryotic replicative CMG helicase. Nat Commun 11, 688 (2020). 10.1038/s41467-020-14577-6

5 Greiwe, J. F. et al. Structural mechanism for the selective phosphorylation of DNA-loaded MCM double hexamers by the Dbf4-dependent kinase. Nat Struct Mol Biol 29, 10–20 (2022). 10.1038/s41594-021-00698-z

6 Pintilie, G. et al. Measurement of atom resolvability in cryo-EM maps with Q-scores. Nat Methods 17, 328–334 (2020). 10.1038/s41592-020-0731-1

7 He, J., Li, T. & Huang, S. Y. Improvement of cryo-EM maps by simultaneous local and non-local deep learning. Nat Commun 14, 3217 (2023). 10.1038/s41467-023-39031-1

8 Croll, T. I. ISOLDE: a physically realistic environment for model building into low-resolution electron-density maps. Acta Crystallogr D Struct Biol 74, 519–530 (2018). 10.1107/S2059798318002425

